# ACE2 is the critical *in vivo* receptor for SARS-CoV-2 in a novel COVID-19 mouse model with TNF- and IFNγ-driven immunopathology

**DOI:** 10.1101/2021.08.09.455606

**Authors:** Riem Gawish, Philipp Starkl, Lisabeth Pimenov, Anastasiya Hladik, Karin Lakovits, Felicitas Oberndorfer, Shane J.F. Cronin, Anna Ohradanova-Repic, Gerald Wirnsberger, Benedikt Agerer, Lukas Endler, Tümay Capraz, Jan W. Perthold, Astrid Hagelkruys, Nuria Montserrat, Ali Mirazimi, Louis Boon, Hannes Stockinger, Andreas Bergthaler, Chris Oostenbrink, Josef M. Penninger, Sylvia Knapp

**Affiliations:** Laboratory of Infection Biology, Department of Medicine I, Medical University of Vienna, Vienna, Austria; Department of Pathology, Medical University of Vienna, Vienna, Austria; Institute of Molecular Biotechnology of the Austrian Academy of Sciences, Vienna, Austria; Molecular Immunology Unit, Institute for Hygiene and Applied Immunology, Center for Pathophysiology, Infectiology and Immunology, Medical University of Vienna, Vienna, Austria; Aperion Biologics, Vienna, Austria; CeMM, Research Center for Molecular Medicine of the Austrian Academy of Sciences, Vienna, Austria; Institute of Molecular Modeling and Simulation, Department of Material Sciences and Process Engineering, University of Natural Resources and Life Sciences, Vienna, Austria; Pluripotency for Organ Regeneration, Institute for Bioengineering of Catalonia (IBEC), The Barcelona Institute of Technology (BIST), Catalan Institution for Research and Advanced Studies (ICREA), Barcelona, Spain; Centro de Investigación Biomédica en Red en Bioingeniería, Biomateriales y Nanomedicina, Madrid, Spain; Karolinska Institute and Karolinska University Hospital, Department of Laboratory Medicine, Unit of Clinical Microbiology, Stockholm, Sweden; National Veterinary Institute, Uppsala, Sweden; Polpharma Biologics, Utrecht, The Netherlands; Department of Medical Genetics, Life Sciences Institute, University of British Columbia, Vancouver, Canada

**Keywords:** mouse-adapted SARS-CoV-2, maVie16, COVID-19 mouse model, C57BL/6, BALB/c, cytokine storm, IFN, TNF, ACE2 targeting, COVID-19 therapy

## Abstract

Despite tremendous progress in the understanding of COVID-19, mechanistic insight into immunological, disease-driving factors remains limited. We generated *maVie16*, a mouse-adapted SARS-CoV-2, by serial passaging of a human isolate. *In silico* modelling revealed how Spike mutations of *maVie16* enhanced interaction with murine ACE2. *MaVie16* induced profound pathology in BALB/c and C57BL/6 mice and the resulting mouse COVID-19 (*mCOVID-19*) replicated critical aspects of human disease, including early lymphopenia, pulmonary immune cell infiltration, pneumonia and specific adaptive immunity. Inhibition of the proinflammatory cytokines IFNγ and TNF substantially reduced immunopathology. Importantly, genetic ACE2-deficiency completely prevented *mCOVID-19* development. Finally, inhalation therapy with recombinant ACE2 fully protected mice from *mCOVID-19*, revealing a novel and efficient treatment. Thus, we here present *maVie16* as a new tool to model COVID-19 for the discovery of new therapies and show that disease severity is determined by cytokine-driven immunopathology and critically dependent on ACE2 *in vivo*.

**Key points:**

- The mouse-adapted SARS-CoV-2 strain *maVie16* causes fatal disease in BALB/c mice and substantial inflammation, pneumonia and immunity in C57BL/6 mice
- TNFα/IFNγ blockade ameliorates *maVie16*-induced immunopathology
- *MaVie16* infection depends on ACE2 and soluble ACE2 inhalation can prevent disease

## Introduction

SARS-CoV-2, was identified as the causative agent of COVID-19 in December 2019 and has since (until August 5^th^, 2021) infected 200 million people and caused 4.3 million confirmed deaths worldwide (WHO, 2021b). Upon adaptation to humans, SARS-CoV-2 evolution gave rise to multiple variants including novel SARS-CoV-2 variants of concern (VOCs), characterized by increased transmissibility and/or virulence, or reduced effectiveness of countermeasures such as vaccinations or antibody therapies (WHO, 2021a). Clinical symptoms upon SARS-CoV-2 infection show a wide range, from asymptomatic disease to critical illness (Chen et al., 2020; Paranjpe et al., 2020). Severe COVID-19 is characterized by progressive respiratory failure, often necessitating hospitalization and mechanical ventilation. Fatal disease is linked to acute respiratory distress syndrome (ARDS), associated with activation of inflammation and thrombosis, often resulting in multi-organ failure (Chen *et al.*, 2020; Paranjpe *et al.*, 2020). Epidemiologically, several risk factors, such as age, male sex, diabetes and obesity have been found to be associated with the development of severe COVID-19 (Richardson et al., 2020; Zhou et al., 2020). However, early immunologic determinants driving disease severity and outcome of SARS-CoV-2 infection remain elusive.

While SARS-CoV-2 replicates primarily in nasopharyngeal and type-II alveolar epithelial cells in the lower respiratory tract, recent data point towards a broader cellular tropism under certain conditions (Sungnak et al., 2020; Ziegler et al., 2020). This largely correlates with angiotensin converting enzyme-2 (ACE2) expression, the entry receptor for SARS-CoV-2 (Hoffmann et al., 2020), and TMPRSS2 expression, a protease, which supports viral cell entry (Hoffmann *et al.*, 2020; Liu et al., 2021; Murgolo et al., 2021). Severe disease seems to be driven by a circuit of T cells and macrophages causing a cytokine release syndrome, immunothrombosis and resulting tissue damage that resembles macrophage activation syndrome (Mangalmurti and Hunter, 2020). As such, IFNγ released by T cells drives the emergence of pathologically activated, hyperinflammatory monocytes and macrophages that produce large amounts of interleukin (IL)-6, TNFα and IL-1β (Grant et al., 2021). Although numerous studies investigated immune responses upon SARS-CoV-2 infection in humans, these studies are largely correlations in hospitalized patients with moderate to severe disease (Merad et al., 2021).

Mechanistic data on early immunologic events driving COVID-19 remain sparse due to limited access to human samples and the scarcity of robust small animal models. Notably, mice are resistant to SARS-CoV-2 infection due to phylogenetic differences in ACE2 (Damas et al., 2020; Parolin et al., 2021). To circumvent the missing sensitivity of conventional mice, some groups studying SARS-CoV-2-related immune mechanisms take advantage of K18-hACE2 transgenic mice, which express human ACE2 under the epithelial cytokeratin-18 (K18) promoter (Winkler et al., 2020). However, K18-hACE2 mice are overly susceptible as they develop deadly SARS-CoV-2 encephalitis (in addition to pneumonia) in response to low infection doses, due to non-physiological and broad (over)expression of hACE2 (Kumari et al., 2021). Recently, four mouse-adapted SARS-CoV-2 strains have been generated by either genetic engineering (Dinnon et al., 2020) and/or serial passaging (Gu et al., 2020; Huang et al., 2021; Leist et al., 2020). However, while infectivity is achieved with all of these strains, only two (Huang et al., 2021; Leist et al., 2020) cause a pathology that mimics human disease, and pathological mechanisms are still poorly understood. Of note, the VOCs Beta/B.1.351 (Tegally et al., 2020) and Gamma/P1 (Voloch et al., 2020) have been recently found to productively replicate in mice without causing pneumonia (Montagutelli et al., 2021), indicating that additional mutations might further increase mouse infectivity and pathogenicity.

In this study, we report the generation of a novel mouse adapted SARS-CoV-2 virus, *maVie16*, which causes a disease reminiscent of COVID-19 in humans in two distinct wild type mouse strains, BALB/c and C57BL/6 mice. We find that the inflammatory cytokines TNF and IFNγ are important drivers of *maVie16* pathogenicity. Moreover, we provide the first genetic *in vivo* proof that ACE2 is the critical SARS-CoV-2 receptor. Finally, therapeutic inhalation of soluble ACE2 completely prevented disease. Our novel mouse *maVie16* infection model is thus a useful and relevant model to study immunology and determinants of SARS-CoV-2 infections and COVID-19 *in vivo*.

## Results

### Generation of the mouse-adapted SARS-CoV-2 strain *maVie16*

To generate a virus pathogenic to mice, we infected BALB/c mice with a human SARS-CoV-2 isolate (BetaCoV/Munich/BavPat1/2020; from here on referred to as *BavPat1* (Rothe et al., 2020)), followed by serial passaging of virus-containing cell-free lung homogenates of infected mice every 3 days (Figure 1A). Infectivity was quickly established after the first few passages, as indicated by increasing SARS-CoV-2 genome copies detected in the lungs of infected mice (Figure 1B). While the viral load did not further change after passage 3, a progressive loss of body weight of mice infected with later stage passaged SARS-CoV-2 indicated enhanced pathogenicity of the virus (Figure 1C). Mice infected with passage 15 furthermore exhibited a severe drop in body temperature (Figure 1D). The transcriptional analysis of pulmonary inflammatory genes (including *il1b*, *ifng*, *il6* and *tnf*) 3 days after infection with the different passages revealed a progressively increasing inflammatory response (Figure 1E). These results indicate that our serial passaging protocol has allowed us to generate a highly infectious and pathogenic mouse-adapted SARS-CoV-2 variant (isolated from lungs of mice infected with passage 15) which we refer to as *maVie16* (**m**ouse-**a**dapted SARS-CoV-2 virus **Vie**nna, passage **16**).

**Figure 1:**
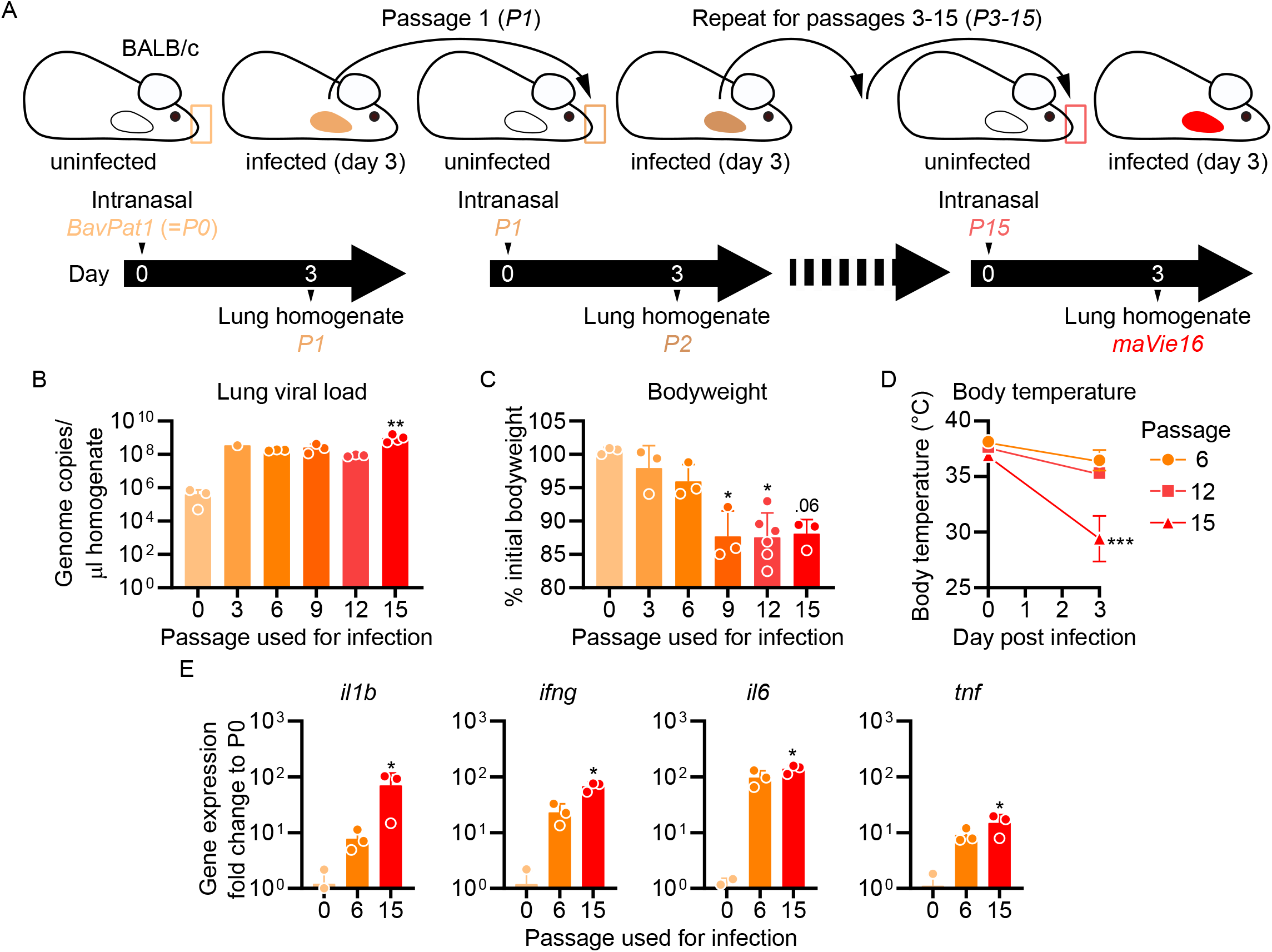
Serial pulmonary passaging of SARS-CoV-2 through BALB/c mice leads to mouse adaptation and generation of the mouse-virulent virus *maVie16*. (A) Experimental strategy for generation of *maVie16*. BALB/c mice were intranasally inoculated with *BavPat1* (Passage 0/P0), followed by serial passaging of virus-containing cell-free lung homogenates of infected mice every 3 days. Passaging was repeated 15 times. (B) Lung tissue virus genome copy numbers (determined by real-time PCR) of mice 3 days after infection with virus of different passages as indicated. (C) Bodyweight (percentage of initial) of mice 3 days after infection. (D) Body temperature before and 3 days after infection. (E) Lung tissue expression fold change (compared to P0 mean; analyzed by real-time PCR) of indicated genes 3 days after infection. (B to E) n = 1-3; (B, C and E): symbols represent individual mice; Kruskal-Wallis test (*vs.* P0) with Dunn’s multiple comparisons test; (D) mean +/−SD; 2way ANOVA with Sidak test (*vs.* the respective initial body temperature); \**P* ≤ 0.05; ** *P* ≤ 0.01; *** *P* ≤ 0.001; numbers above bars show the actual *P* value.

### Genetic evolution of the *maVie16* Spike protein

To elucidate the mechanisms of mouse adaptation of *maVie16*, we sequenced SARS-CoV-2 genomes at all passages and compared the viral sequences to the original *BavPat1* isolate. Despite the substantially increased virulence in mice, we found only a limited number of mutations with high (> 0.5) allele frequencies, located in the Spike protein, open reading frame (ORF)1AB, and envelope (Figure 2A). Given the importance of Spike for SARS-CoV-2 infectivity, we were particularly interested in the three *de novo* mutations within its receptor binding domain (RBD) (Figure 2B). Most prominently, we observed the immediate appearance of Q498H after passage 1 (Figure 2B), which correlated with the early increase in viral load (and hence viral propagation) in infected mice (Figure 1B). After passage 12, we detected a glutamine to arginine exchange at position Q493R (Figure 2B). As of today, Q498H or Q493R have been observed in a few patient samples (16 and 196, respectively, according to the GSAID database (Shu and McCauley, 2017)) whereas the simultaneous appearance of both mutations was never reported in human isolates. Q498H and Q493R appear to have emerged sporadically and have not yet been reported in specific human SARS-CoV-2 strains. However, both mutations were reported in other mouse-adapted virus strains (Dinnon *et al.*, 2020; Gu *et al.*, 2020; Huang *et al.*, 2021), supporting their critical role in the increased mouse specificity. In contrast to these apparently mouse-specific Spike RBD mutations, the K417 mutation (which appeared around passage 10; Figure 2B) has also been reported in two human SARS-CoV-2 variants of concern, i.e. Beta/B.1.351 (K417N) (Tegally *et al.*, 2020) and Gamma/P1 (K417T) (Voloch *et al.*, 2020).

**Figure 2:**
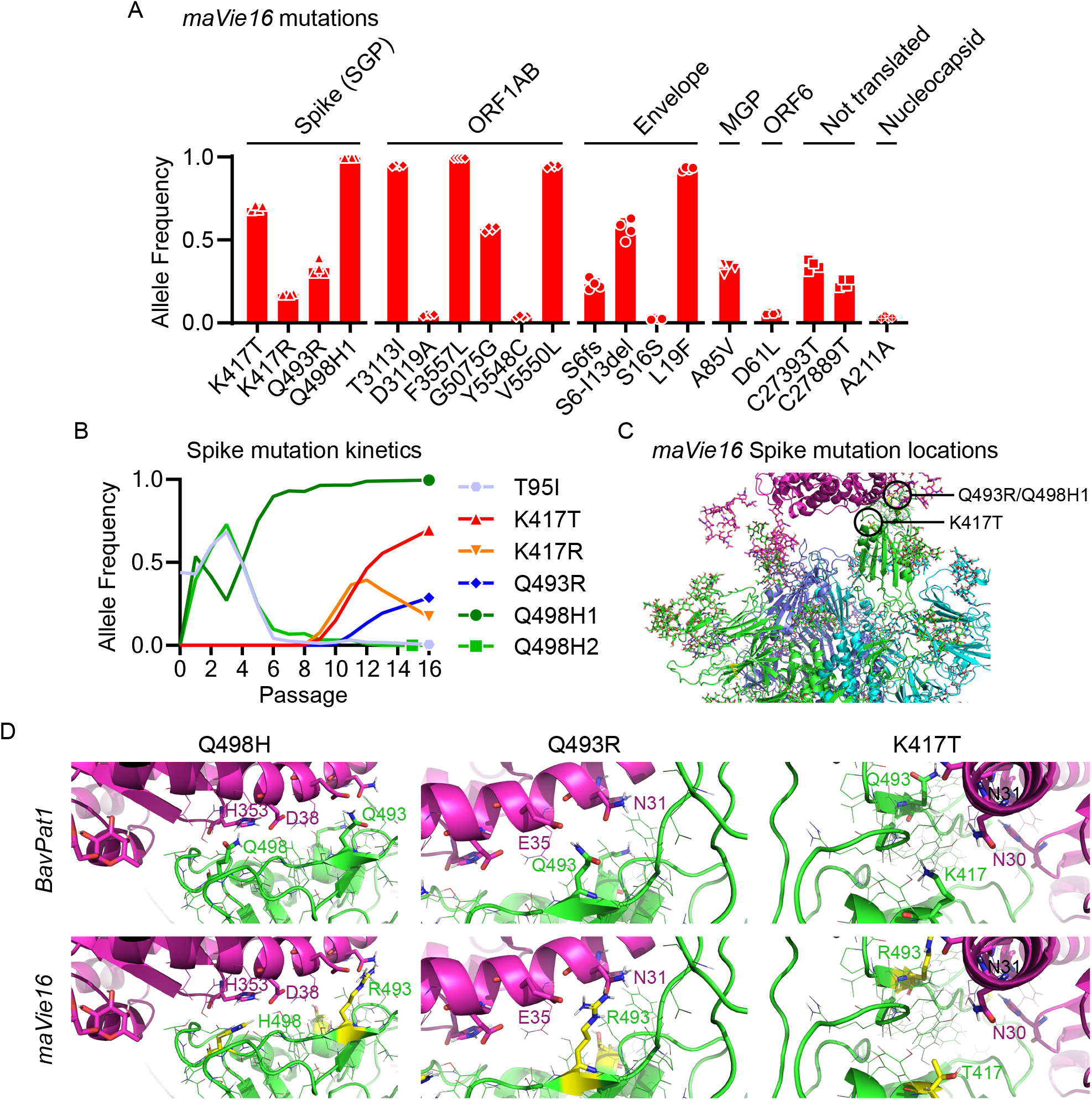
*maVie16* possesses a distinct pattern of mutations and mediates *in vivo* pathology via ACE2. (A) Overview of allele frequencies of mutated amino acids detected in *maVie16* by sequencing. Labels on top indicate the associated protein (SGP: Spike glycoprotein; ORF: open reading frame; MGP: membrane glycoprotein). (B) Spike protein mutation dynamics. (C) Modelling and location of spike mutations. Spike trimer in cyan blue and green, mACE2 in magenta cartoon representation. Glycans in stick representations. (D) Modelling of specific *BavPat1* (upper row) and *maVie16* (lower row) amino acid regions in green (respective mutated positions are highlighted in yellow and labelled in green) and their interaction with mouse ACE2 (in magenta; positions of interest are labelled in magenta or black).

Structural analyses of mouse and human ACE2 (mACE2 and hACE2, respectively) have shown that the surface of mACE2 consists of more negative charged amino acids than that of hACE2 (Figure S1A) (Rodrigues et al., 2020). Indeed, the exchange of two glutamines (Q493 and Q498) to arginine and histidine, respectively, in the *maVie16* Spike results in a positive charged surface (Figure 2C). Detailed structural modeling of the *BavPat1 versus* the *maVie16* Spike protein interface with mACE2 revealed that the *maVie16* Spike Q498H mutation would strengthen its interaction with mACE2 at amino acid D38, otherwise forming an intramolecular salt bridge with mACE2 H353 (Figure 2D). Similarly, the newly-introduced arginine at *maVie16* spike position 493 is predicated to efficiently interact with the negatively-charged mACE2 amino acid E35, thus further stabilizing the *maVie16* Spike/mACE2 interaction by neutralizing the otherwise unpaired negative charge of mACE2 glutamic acid (Figure 2D). Finally, the threonine at Spike position 417 (T417) of *maVie16*’s is predicted to enhance the interaction with the neutrally-charged asparagine at position 30 of mACE2 while the aspartic acid at this position in hACE2 forms a salt-bridge with lysine K417 of the original *BavPat1* isolate. Of note, *maVie16* showed similar propagation kinetics in Vero and Caco-2 cells as compared to *BavPat1* (Figure S1B). Thus, while the interaction with mACE2 is enhanced, the infectivity of human cells remained unchanged for *maVie16* and both simian and human cells continue to serve as a useful tool for *maVie16* propagation and titer determination. Taken together, our data indicate that only three distinct spike RBD mutations introduced during passaging could substantially enhance interaction with mACE2, without obvious effects on the interaction with hACE2.

### *MaVie16* causes severe pneumonia in BALB/c and C57BL/6 mice

We next performed dose response experiments of *maVie16* infections in BALB/c (B/c) and, as a commonly used mouse strain for genetic engineering, C57BL/6 (B/6) mice and monitored weight and body temperatures over 7 days. In BALB/c mice, *maVie16* was highly pathogenic and caused profound weight loss, starting around day 3 post infection (p.i.) at a dose of 4 × 10^3^ TCID_50_ (Figure 3A), whereas all tested higher doses caused an earlier weight loss (day 2 p.i.) and lethality starting at day 4 p.i. (Figure 3B). Notably, the loss of body weight correlated with hypothermia (Figure 3C).

**Figure 3:**
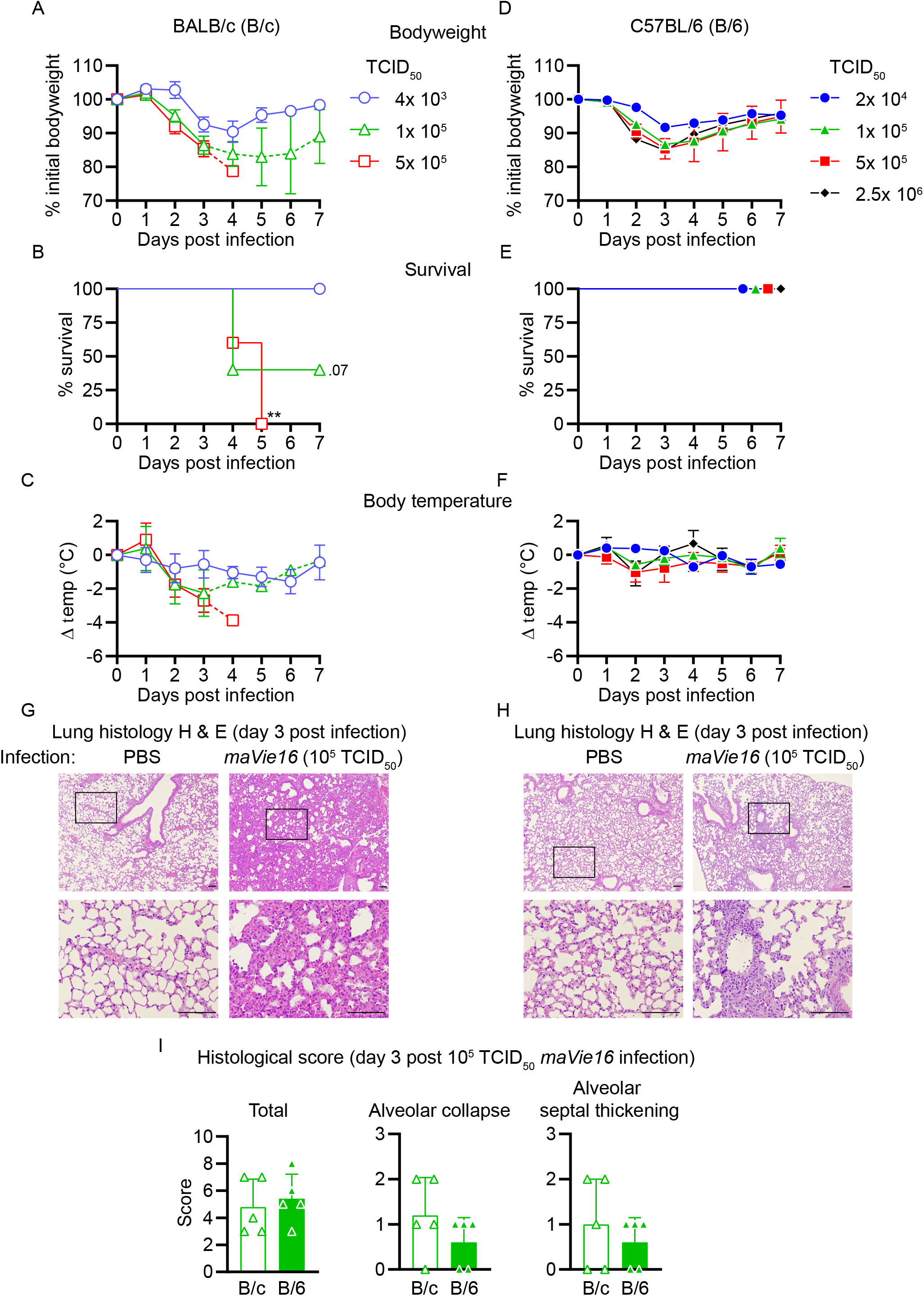
Respiratory *maVie16* infection causes dose-dependent pathology in BALB/c and C57BL/6 mice. (A-C and G) BALB/c (B/c) or (D-F and H) C57BL/6 (B/6) mice were intranasally inoculated with different doses of *maVie16* as indicated and monitored for (A and D) bodyweight, (B and E) survival and (C and F) body temperature over 7 days; dashed lines in A and C indicate trajectories of groups lacking full group size due to death of animals (see B). (G and H) Lung sections (hematoxylin & eosin stain) from mice three days after infection with 10^5^ TCID_50_ *maVie16*; black rectangles in the upper pictures indicate the area magnified in the respective lower row picture; scale bars indicate 100 μm. (I) Histological score for analysis of lung sections as described in (G) and (H); symbols represent individual mice; (A, C, D, F) mean +/−SD; (B) Mantel-Cox test (*vs.* 4x 10^3^ TCID_50_); ** *P* ≤ 0.01; the number next to the symbol shows the actual *P* value;

In contrast to BALB/c mice, *maVie16* infection did not cause lethality nor significant hypothermia in C57BL/6 animals, albeit 1 × 10^5^ (and higher) TCID_50_ induced a profound but transient body weight loss of about 15-20% (Figure 3D to F). Interestingly, both mouse strains infected with 1 × 10^5^ TCID_50_ showed similar body weight loss (approximately 15%) at day 3 p.i., but while disease severity further increased in BALB/c animals, C57BL/6 mice recovered. In line with these data, BALB/c animals showed more severe lung pathologies compared to C57BL/6 mice, as revealed by histologic analyses of lungs on day 3 (Figure 3G and H). While the cumulative histological score revealed no differences between the strains, diffuse alveolar damage, characterized by alveolar collapse and septal thickening, was more pronounced in BALB/c mice (Figure 3I).

To compare our data with a widely used mouse model of COVID-19, we infected *K18-hACE2* transgenic mice (which express human ACE2 under control of the human keratin 18 promoter (McCray et al., 2007; Oladunni et al., 2020) with either 1 × 10^3^ or 1 × 10^4^ TCID_50_ of *BavPat1* and monitored the disease course over 7 days. As expected (Oladunni *et al.*, 2020), *K18-hACE2* mice were most sensitive to infection and succumbed to *BavPat1* infections (Figure S2A and S2B). However, the disease course was profoundly different to that of *maVie16*-infected wild type animals. *BavPat1*-infected *K18-hACE2* mice appeared healthy until day 4 p.i. and then progressively lost weight until day 7, accompanied by hypothermia and death (Figure S2C-F). Death in *K18-hACE2* mice most likely results from severe encephalitis due to expression of hACE2 in neurons where ACE2 is normally not expressed (Hikmet et al., 2020). These results show that *maVie16* causes severe pathology in BALB/c and profound, but transient, disease in C57BL/6 mice, recapitulating critical aspects (such as transient pneumonia and viral clearance) of human COVID-19.

### Murine anti-SARS-CoV-2 immune responses

To better understand the pathophysiology of *mCOVID-19*, we next profiled immune cell dynamics in blood and lungs of C57BL/6 mice during acute infection and upon recovery. *MaVie16* (5 × 10^5^ TCID_50_) infection caused severe blood leukopenia on day 2 p.i. with a prominent reduction of lymphocytes, monocytes, neutrophils and NK cells (Figure 4A, Figure S3 and S4A). At the same time, we observed an expansion of peripheral plasmacytoid dendritic cells (pDCs) (Figure 4B), which are important antiviral effector cells capable of efficient and rapid type I interferon (IFN) production (Swiecki and Colonna, 2015).

**Figure 4:**
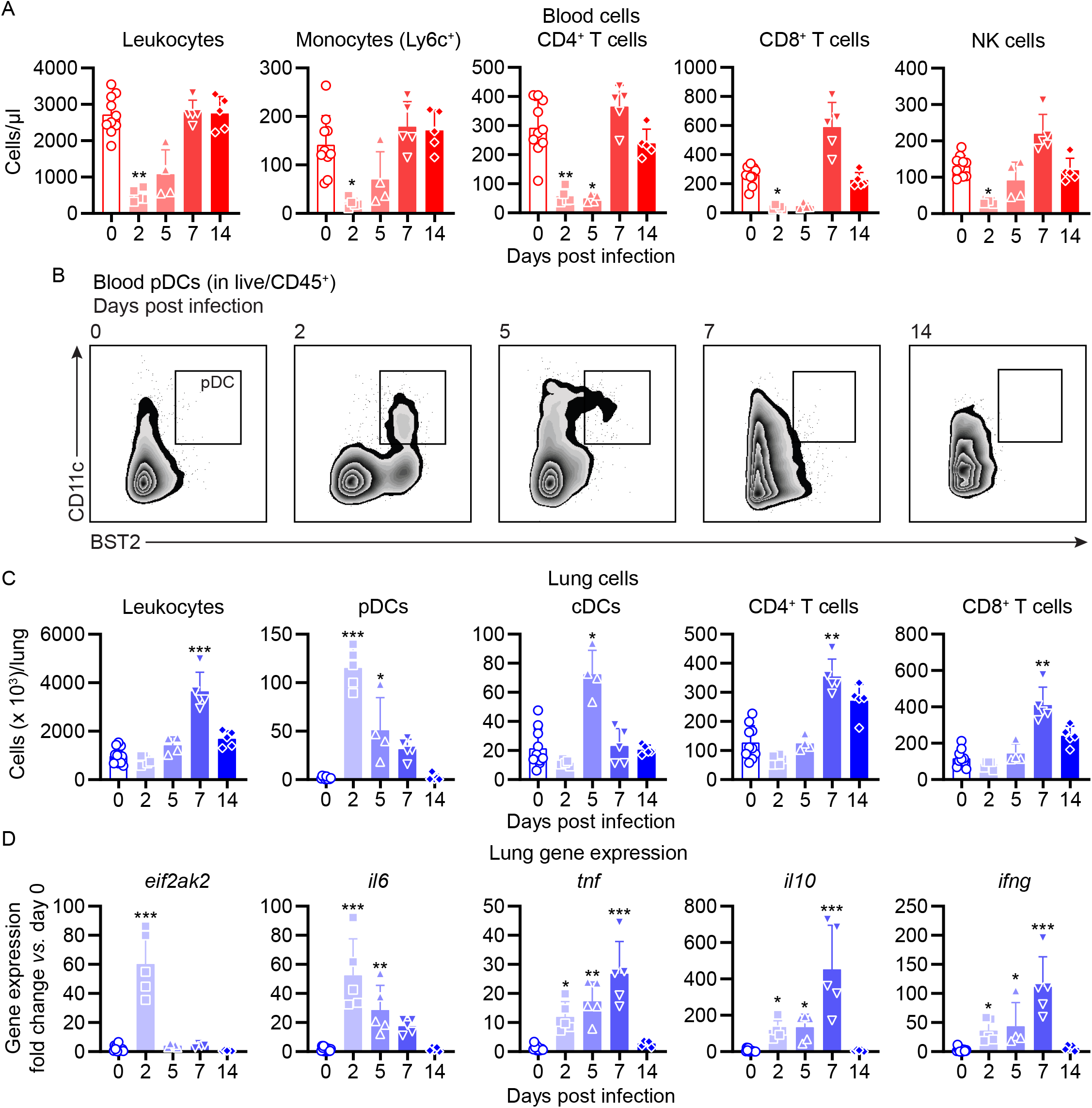
*mCOVID-19* is associated with transient lymphopenia, pulmonary dendritic cell and T cell infiltration and pneumonia. C57BL/6 mice were intranasally infected with PBS (= group 0) or 5×10^5^ TCID_50_ *maVie16* and sacrificed after 2, 5, 7 or 14 days for subsequent analysis. (A) Flow cytometry analysis of blood cell populations. (B) Density plot representation of blood plasmacytoid dendritic cells (pDCs; identified as live/CD45^+^/CD11c^+^/BST2^+^) analyzed by flow cytometry. (C) Flow cytometry analysis of whole lung cell populations (see Figure S3 for gating strategies). (D) Lung tissue expression fold change (compared to group 0 mean; analyzed by real-time PCR) of indicated genes from mice at the respective time points after infection. (A, C and D): symbols represent individual mice; Kruskal-Wallis test (*vs.* group 0) with Dunn’s multiple comparisons test; \**P* ≤ 0.05; ** *P* ≤ 0.01; *** *P* ≤ 0.001.

Infection led to substantial pulmonary infiltration with leukocytes (Figure 4C), paralleling the development of pneumonia. A closer look at the inflammatory cell composition in the lungs revealed a remarkable infiltration with pDCs on day 2 p.i. (Figure 4C). Conventional dendritic cells (cDCs) transiently accumulated in the lung on day 5 p.i., followed by T helper and cytotoxic T cell infiltration and recruitment of monocytes on day 7 (Figure 4C and S4B). The abundance of other analyzed immune cell populations did not change substantially, and lung neutrophil numbers dropped around day 2 p.i. to return to baseline levels over the remaining disease course (Figure S4B). Surprisingly, pulmonary NK cells, which fulfill important antiviral functions (Bjorkstrom et al., 2021), did not significantly expand in the lungs of *maVie16* infected C57BL6 mice (Figure S4B). In line with an early lung pDC accumulation, we found rapid induction of type I IFN-inducible genes, such as *eif2ak2* (protein kinase R/PKR) and *ifit1*, accompanied by a profound *il6* response; the expression of *tnf*, *il10*, *il1b* and *tgfb* peaked between day 5 and 7 p.i. (Figure 4D and S4C). The *ifng* induction reached a maximum on day 7, coinciding with the increased numbers of T cells which are prominent sources of IFNγ (Figure 4D).

In C57BL/6 mice infected with *maVie16* (5 × 10^5^ TCID_50_), lung weights significantly increased by day 5 to 7 p.i. (Figure 5A). Lung viral titers peaked at day 2 p.i. and then gradually declined (Figure 5B). Immunohistochemical staining for SARS-CoV-2 nucleoprotein on lung slides confirmed the initial presence and subsequent elimination of viral particles (Figure 5C). Histological analyses of lung tissue revealed progressive interstitial and perivascular infiltration and signs of diffuse alveolar damage, such as alveolar collapse and septal thickening, peaking from day 5 to 7, and resolving by day 14 p.i. (Figure 5C). Interestingly, vasculitis was observed early after infection (day 2) and resolved at later timepoints (Figure 5C and 5D). Moreover, most acute inflammatory parameters (including cytokine genes, innate immune cells and lung tissue weights) had returned close to baseline at day 14 p.i. (Figures 4, 5A-D and S4), confirming resolution of inflammation. Onset of adaptive immunity, as indicated by expansion of pulmonary cDCs and T helper cells by day 5 and 7 (Figure 4C), as well as by increased spleen weight on day 7 p.i. (Figure S4D) was followed by an efficient humoral immune response, reflected by significantly increased levels of anti-SARS-CoV-2 Spike protein-specific plasma IgG1, IgG2b and IgA antibodies on day 14 p.i. (Figure 5E). These data show that *maVie16* infection causes severe, yet transient, lung inflammation in C57BL/6 mice. Furthermore, *maVie16* induces a rapid type I IFN response, which correlated with a marked expansion of pDCs.

**Figure 5:**
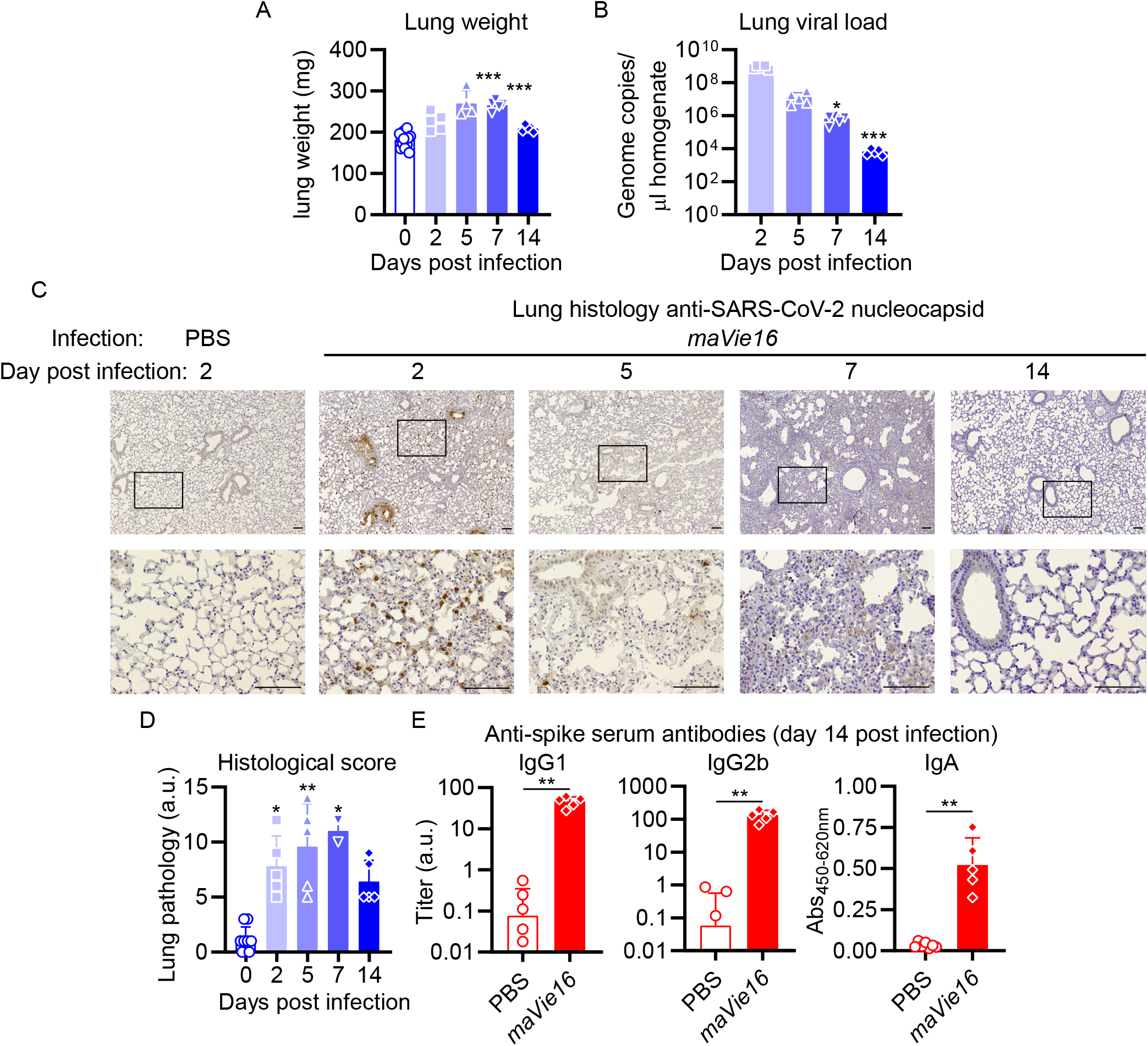
*mCOVID-19* induces transient pneumonia and antigen-specific adaptive immunity. C57BL/6 mice were intranasally infected with PBS (= group 0 or PBS) or 5x 10^5^ TCID_50_ *maVie16* and sacrificed after 2, 5, 7 or 14 days for subsequent analysis. (A) Lung tissue virus genome copy numbers (determined by real-time PCR). (B) Lung tissue weight. (C) Representative lung immunohistochemistry (anti-SARS-CoV-2 nucleocapsid stain, counterstained with hematoxylin) pictures; black rectangles in the upper pictures indicate the area magnified in the respective lower row picture; scale bars represent 100 μm. (D) Lung pathology score based on histological analysis of lung tissue sections. (E) Analysis (by ELISA) of SARS-CoV-2 spike-specific IgG1, IgG2b and IgA plasma antibody titers 14 days after infection. (A, B, D, E) mean +SD; symbols represent individual mice; (A, B, D): Kruskal-Wallis test (*vs.* (A) day 2 or (B and D) group 0) with Dunn’s multiple comparisons test; (E) Mann-Whitney test; \**P* ≤ 0.05; ** *P* ≤ 0.01; *** *P* ≤ 0.001;

### Distinct differences in anti-viral immunity of BALB/c *vs.* C57BL/6 mice

In an attempt to elucidate underlying mechanisms for the different susceptibility of BALB/c and C57BL/6 mice, we infected both mouse strains side by side with 1 × 10^5^ TCID_50_ of *maVie16* and monitored their respective immune responses 3 days later (as BALB/c and C57BL/6 mice showed comparable weight loss and no mortality at this dose and time point; Figure 3). Both mouse strains exhibited a decline in blood leukocytes, including lymphocytes and NK cells, upon infections with *maVie16* (Figure 6A and S5A). Also, pulmonary immune cell dynamics did not differ between the strains, except for an expansion of NK cells in BALB/c mice, and elevated pDC numbers in C57BL/6 animals (Figure 6B and S5B). Higher NK cell abundance correlated with increased pulmonary IFNγ levels in BALB/c mice, whereas *Il1b* was slightly higher in C57BL/6 animals 3 days after infection (Figure 6C and S5C). Plasma cytokine levels (Figure S5D), viral loads (Figures S5E) as well as spleen and lung weight (Figure S5F) were comparable between the two mouse strains. These data identify differences in the early local inflammatory response between *maVie16*-infected BALB/c as compared to C57BL/6 animals.

**Figure 6:**
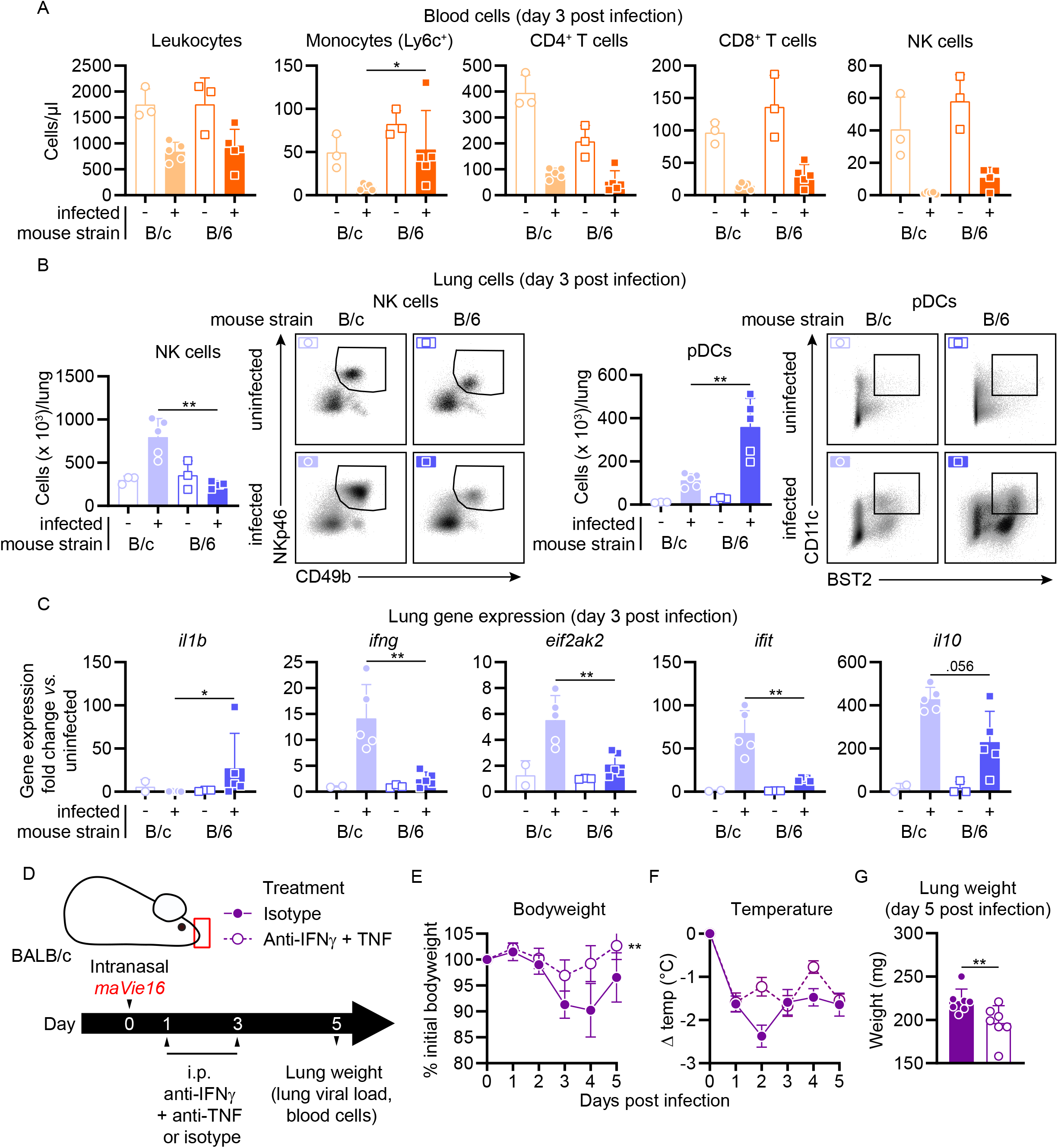
BALB/c *mCOVID-19* is associated with an increased NK cell and interferon response and is ameliorated by IFNγ and TNF blockade. BALB/c (B/c) and C57BL/6 (B/6) mice were intranasally inoculated with 10^5^ TCID_50_ *maVie16* (+) or PBS (−). Samples for analyses were collected 3 days after infection. (A) Flow cytometry analysis of blood cell populations. (B) Flow cytometry analysis of whole lung NK cells and plasmacytoid dendritic cells (pDCs). Density plots represent examples of respective cell populations (NK cells pre-gated from live/CD45^+^/Ly6G^−^/CD3^−^; pDCs pre-gated from live/CD45^+^) (C) Lung tissue expression fold change (compared to the respective mean of uninfected samples; analyzed by real-time PCR) of indicated genes. (D) Experimental scheme for E to G. BALB/c mice were infected with 10^5^ TCID_50_ *maVie16* and treated intraperitoneally on day 1 and 3 p.i. with a mix of 500 μg anti-IFNγ and anti-TNF or with isotype control antibody. (E) Bodyweight and (F) temperature kinetics over 5 days after infection. (G) Lung weight on day 5 after infection. (A to C, G) mean +SD; symbols represent individual mice; differences between infected groups were assessed using the Mann-Whitney test; (E and F) mean +/−SD; 2way ANOVA with Dunnett’s multiple comparisons test (*vs.* the respective initial bodyweight or temperature); in panels without respective labels, the groups were not significantly different (*P* < 0.05); \**P* ≤ 0.05; ** *P* ≤ 0.01;

### A pathogenic role of IFNγ and TNF in disease severity

Several reports illustrated a correlation between systemic cytokine responses, lung injury and prognosis in humans suffering from severe COVID-19. A particularly detrimental role was attributed to excessive IFNγ levels (Grant *et al.*, 2021). Moreover, an earlier study showed that the combined activity of IFNγ and TNF caused inflammatory types of cell death, which was named PANoptosis, i.e. pyroptosis, apoptosis and necroptosis, and that blocking these cytokines improved disease outcomes in *K18-hACE2* mice infected with human SARS-CoV-2 (Karki et al., 2021). In accordance with these reports and our finding of higher IFNγ levels in highly susceptible BALB/c mice, we tested if blocking IFNγ and TNF might reduce disease severity after *maVie16* infection. *MaVie16*-infected (1 × 10^5^ TCID_50_) BALB/c mice treated intraperitoneally with a mixture of anti-IFNγ and anti-TNF antibodies (on days 1 and 3 p.i.; Figure 6D and S6A) significantly improved weight loss (Figure 6E) and decreased lung weights (Figure 6G), the viral load (Figure S6B) as well as circulating monocyte levels (Figure S6C) while it had only minor effects on body temperature (Figure 6F). In C57BL/6 mice, the same treatment had no effects on body or lung weights, or viral load (Figure S6D, E, G and H), but significantly improved the altered body temperature and circulating leukocyte numbers (Figure S6F and I). Though other cytokines and chemokines must be involved, these results underline the therapeutic potential of IFNγ and TNF blockade as COVID-19 treatment.

### ACE2 expression is essential for *maVie16* infections

We have previously shown using *Ace2* mutant mice that ACE2 is essential for *in vivo* SARS infections (Kuba et al., 2005). ACE2 is also an important receptor for SARS-CoV-2, however other receptors have been critically proposed to mediate infections such as neuropilin-1 (Cantuti-Castelvetri et al., 2020; Daly et al., 2020) or CD147 (Wang et al., 2020). Having developed the *maVie16* infection system, we could therefore ask one of the key questions to understand COVID-19: is ACE2 also the essential entry receptor for SARS-CoV-2 infections in a true *in vivo* infection model? To answer this question, we infected ACE2-deficient male *Ace2*^-/*y*^ mice (Figure 7A and S7A). In contrast to infected littermate control animals, *Ace2*^-/*y*^ mice were fully protected against (5× 10^5^ TCID_50_) *maVie16*-induced pathology and showed stable body weight (Figure 7B) and temperature (Figure 7C). By day 3 p.i., ACE2 deficiency protected from pneumonia development as indicated by lower lung weight (Figure 7D) and the absence of any lung pathology (Figure 7E). Viral genome copy numbers at this time point were similar (Figure S7B). However, while infected cells were evenly distributed in wild type control lungs, we did not find any SARS-CoV-2 nucleocapsid-positive cells in lungs of *Ace2*^-/*y*^ animals (Figure 7F). These data support a gatekeeper role for ACE2 in mediating productive infection of lung epithelial cells with *SARS-CoV-2.* Thus, genetic inactivation of ACE2 protects mice from productive SARS-CoV-2 infections and lung pathologies.

**Figure 7:**
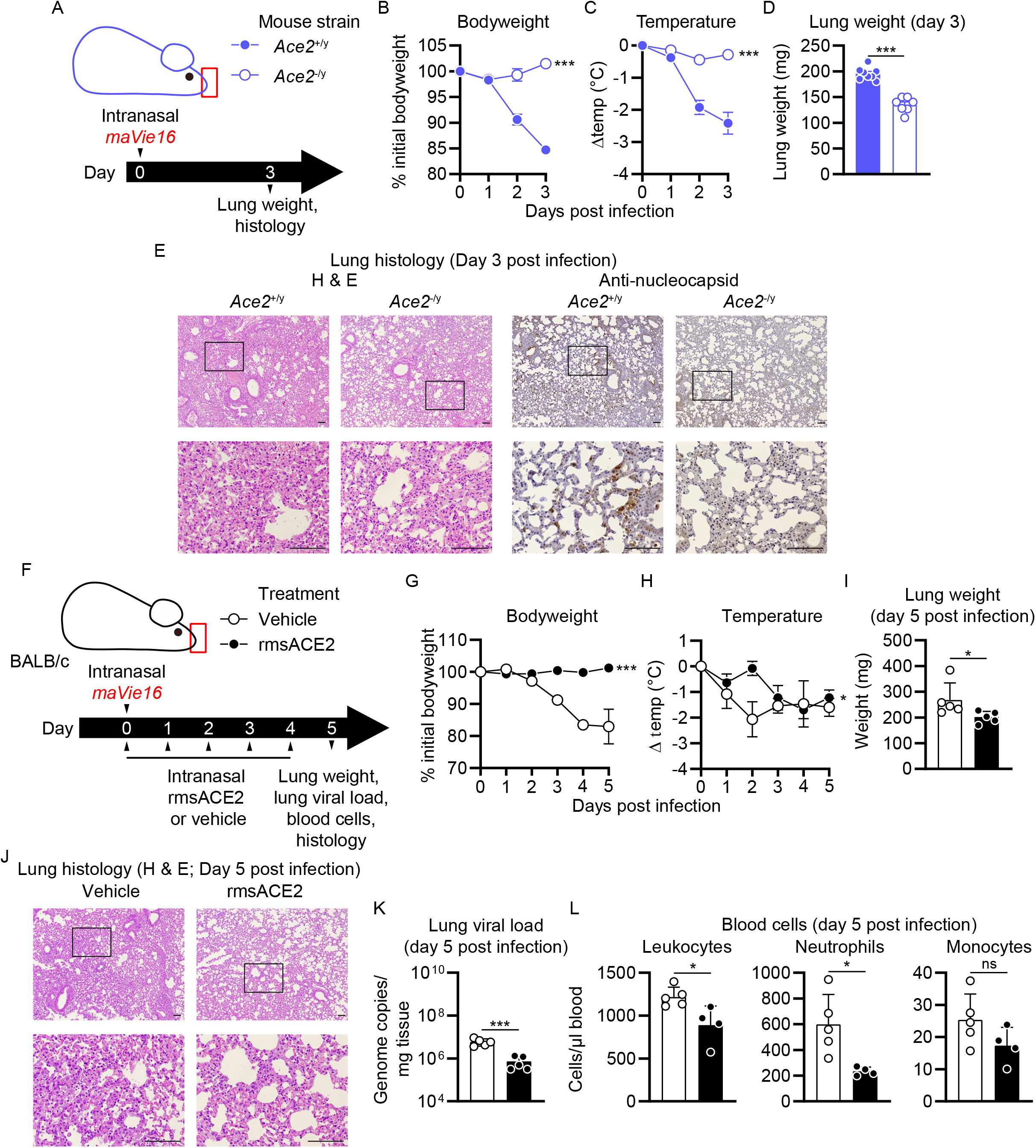
*mCOVID-19* pathology depends on *Ace2* and is improved by recombinant ACE2 administration. (A) Experimental scheme for B to E: Male *Ace2*-deficient (*Ace2*^−/*y*^) or control (*Ace2*^+/*y*^) mice were infected with 5x 10^5^ TCID_50_ *maVie16*. (B) Bodyweight and (C) temperature kinetics over 3 days after infection. (D) Lung tissue weight 3 days post infection (p.i.). (E) Lung histology 3 days after infection (left panels: hematoxylin & eosin stain; right panels: anti-SARS-CoV-2 nucleocapsid immune-stain); black rectangles in the upper pictures indicate the area magnified in the respective lower row picture; scale bars represent 100 μm. (F) Experimental scheme for G to L. BALB/c mice were infected with 10^5^ TCID_50_ *maVie16* and treated daily intranasally up to day 4 p.i. with 100 μg recombinant murine soluble (rms) ACE2 or vehicle (the first treatment was administered together with virus). (G) Bodyweight and (H) temperature kinetics over 5 days after infection. (I) Lung weight, (J) lung histology (hematoxylin & eosin stain), (K) lung viral load, and (L) blood cells on day 5 after infection. (B, C, G, H) mean +/−SD; 2way ANOVA with Dunnett’s multiple comparisons test (*vs.* the respective initial bodyweight or temperature); (D, I, K, L) Mann-Whitney test; * *P* ≤ 0.05; ** *P* ≤ 0.01; *** *P* ≤ 0.001; ns: not significant (*P* > 0.05)

### Inhalation of ACE2 can protect from *maVie16* infections

To test whether ACE2 could also be used therapeutically to protect from *maVie16*-induced lung pathologies in BALB/c and C57BL/6 mice, we performed therapeutic inhalation treatments with recombinant murine soluble (rms) ACE2. *MaVie16*-infected (1 × 10^5^ TCID_50_) BALB/c mice which were treated with rmsACE2 (Figure 7F and S7C), did not lose body weight (Figure 7G), developed less hypothermia (Figure 7H), exhibited lower lung weights (Figure 7I) and were largely protected from pneumonia (day 5 p.i.; Figure 7J), had lower lung viral loads (Figure 7K) and fewer blood neutrophils (Figure 7L) than vehicle-treated animals, indicating marked prevention of *mCOVID-19* development. Further, we found fewer SARS-CoV-2 nucleocapsid-positive cells in lungs of BALB/c animals treated with rmsACE2 5 days after infection with *maVie16* as compared to controls (Figure S7D). Similarly, rmsACE2 treatment of *maVie16*-infected (5 × 10^5^ TCID_50_) C57BL/6 mice (Figure S7E) prevented weight loss and reduced the viral load (Figure S7F and G, respectively) while it did not affect temperature, lung weight and blood leukocyte numbers (Figure S7G, H and J, respectively). These data show that inhalation of rmsACE2 can markedly prevent *maVie16*-induced (m)COVID-19.

## Discussion

With *maVie16*, we here generated a novel tool to study and experimentally dissect COVID-19 in vivo. Our passaging protocol led to an initial fast adaptation of the *BavPat1* to the new host, indicated by a rapid increase in virus genome copy number found in the lung. This increased mouse infectivity coincided with the appearance of a spike protein mutation at position 498, which was intentionally introduced using an engineering approach by the Baric group in an earlier attempt to increase mouse specificity of SARS-CoV-2 (Dinnon *et al.*, 2020). Interestingly, another *maVie16* spike protein mutation (at position 493) was also found in the mouse adapted variant of the engineered virus after 10 passages through BALB/c mice (Gu *et al.*, 2020; Huang *et al.*, 2021; Leist *et al.*, 2020) and reported to improve spike-mAce2 interaction (Adams et al., 2021).

The K417T Spike protein mutation was also found in the Gamma/P1 SARS-CoV-2 VOC (Voloch *et al.*, 2020) and the same lysine was changed to an asparagine in the Beta/B.1.351 VOC (Tegally *et al.*, 2020). Both variants have been shown to replicate in laboratory mice (Montagutelli *et al.*, 2021) and in both K417T/N was exclusively observed in conjunction with the mutations N501Y and E484K. While N501Y is suspected to boost infectivity, E484K might contribute to antibody escape and immune evasion. In its interaction with hACE2, E484 is postulated to form a salt-bridge with K31 in hACE2, while the neighboring D30 interacts with K417. Mutating K417 and E484 leads to an intramolecular salt-bridge between D30 and K31 in hACE2, having only minor effects on receptor interaction (Cheng et al., 2021). As outlined above, with aspartic acid replaced by asparagine at position 30 in mACE2, spike protein amino acid K417 no longer has a strong interaction partner and mutation to T417 in *maVie16* is favorable. These data indicate that K417T *per se* might not itself be involved in immune escape mechanisms, but rather emerged to stabilize the interaction with mACE2. Our *in silico* modeling provides strong evidence that these spike protein mutations in *maVie16* support the molecular interaction with mACE2, thereby facilitating infections and the development of severe COVID-19 in mice. At the same time, structural analysis did not support an altered binding of *maVie16* Spike to hACE2, which was confirmed by propagation experiments with *BavPat1* and *maVie16* in human Caco-2 cells.

While inducing profound disease and mortality in BALB/c mice, *maVie16* also induced substantial bodyweight loss and pneumonia in adult C57BL/6 mice. In addition, C57BL/6 mCOVID-19 recapitulated critical immunological aspects observed in human COVID-19, including peripheral leuko- and lymphopenia, pneumonia and antigen-specific adaptive immunity. In addition, mCOVID-19 is characterized by an early pDC mobilization in blood and lung, associated with a fast pulmonary type I IFN response. The susceptibility of C57BL/6 animals proves particularly valuable since a large number of genetically modified mice is available on this background. A side-by-side comparison of BALB/c versus C57BL/6 mice revealed increased lung expression of type II IFN and IFN-driven genes in the BALB/c background, supporting the notion that pronounced inflammation is linked to more severe pathology. A detrimental role for cytokines, specifically IFNγ and TNF, had been previously associated with severe COVID-19 and experimentally identified as risk factor for severe disease in SARS-CoV-2-infected *K18-hACE2* mice, by causing a form of inflammatory cell death termed *PANoptosis* (Karki *et al.*, 2021). We also tested this hypothesis, and found that administration of IFNγ- and TNF-blocking antibodies significantly reduced disease burden and inflammation in *maVie16*-infected BALB/c mice. The role of other cytokines and chemokines as well as innate antiviral immunity or defined cell populations can now be genetically and experimentally dissected using our *maVie16* infection model.

Several studies based on crystallography (Wrapp et al., 2020; Yan et al., 2020), modeling (Rodrigues *et al.*, 2020) or *in vitro* experiments (Cai et al., 2021; Hoffmann *et al.*, 2020; Monteil et al., 2020) propose ACE2 as the main entry receptor for SARS-CoV-2. The relevance of ACE2 is further supported by *in vivo* experiments using *K18-hACE2* mice (Oladunni *et al.*, 2020) or viral vector-mediated hACE2 delivery systems (Rathnasinghe et al., 2020). Although these experiments have shown that ACE2 expression is sufficient for infection, definitive proof of an essential *in vivo* role of ACE2 in COVID-19 has not been provided so far. This is of particular importance because additional receptors have been proposed that could also permit SARS-CoV2 infections (Cantuti-Castelvetri *et al.*, 2020; Daly *et al.*, 2020; Wang *et al.*, 2020). Using *Ace2*-deficient mice, we here demonstrate the strict dependence of *maVie16* infectivity and pathogenicity on ACE2 expression *in vivo*. Furthermore, we exploited this mechanistic insight and treated animals with rmsACE2 inhalation and observed a substantial protection of BALB/c and C57BL/6 mice from developing *mCOVID-19*. A recently published genome-wide association study across more than 50 000 COVID-19 patients and more than 700 000 non-infected individuals revealed a rare variant that was associated with down-regulated *ACE2* expression and reduced risk of COVID-19 (Horowitz *et al.*, 2021), providing additional evidence for the importance of ACE2 for SARS-CoV-2 infection in humans. The principal effectiveness of recombinant human soluble (rhs) ACE2 towards SARS-CoV-2 was demonstrated earlier in organoids, as it slowed viral replication in this setting (Monteil *et al.*, 2020). It has been recently proposed that soluble ACE2 might enhance SARS-CoV-2 infections (Yeung et al., 2021), however our data do not support this. Moreover, using hamsters (Higuchi et al., 2021; Linsky et al., 2020) and the *K18 hACE2* mouse model (Hassler et al., 2021), it has also been shown that different forms of ACE2 can protect from disease, providing – together with our novel *murine COVID-19* model – pre-clinical proof-of-concept that ACE2 can be used as a therapy. RhsACE2, prepared in the same way as rmsACE2 we used for our inhalation experiments (Monteil *et al.*, 2020), is now being tested in clinical trials in COVID-19 patients with first promising results (Zoufaly et al., 2020). Considering the emergence of variants that can escape in part vaccine efficacy and approved antibody therapies (Yuan et al., 2021; Zhou et al., 2021) and the fact that these variants have been apparently selected for better ACE2 binding (Tian et al., 2021; Yuan *et al.*, 2021), ACE2 inhalation has the potential to prevent or treat early stages of SARS-CoV-2 infections irrespective of the virus variant. However, whether inhalable ACE2 indeed constitutes a universal strategy against current and future SARS-CoV-2 variants needs careful testing in clinical trials.

Overall, we here report the development of a novel mouse-adapted SARS-CoV-2, which induces *mCOVID-19* in both BALB/c and C57BL/6 backgrounds. Our findings on the essential role of ACE2 in *in vivo* infections and of recombinant mACE2 administration and IFNγ/TNF blocking as therapeutic options provide a first glimpse of the potential of this new tool to increase our understanding of COVID-19 *in vivo* of to foster the discovery of novel therapeutic options.

## Author contributions

R.G., P.S., L.P., and A.Hl. performed all of the experiments involving SARS-CoV-2, including isolation, culture and animal experiments. K.L and L.P. performed PCR experiments, A.Hl. performed all histological staining. F.O. analyzed lung histology slides. A.R. generated TMPRSS2 overexpressing Vero cells, supported by H.S., G.W. produced and provided mrACE2. T.C., J.W.P. and C.O. performed the molecular modeling, and B.A., L.E. and A.B. sequenced and analyzed viral strains. L.B. provided critical reagents. S.J.F.C., A.Ha. and J.M.P. provided critical reagents and contributed to experimental design of ACE2-related studies. N.M. and A. M. provided critical input on experiments and supported infectious research. R.G., P.S. and S.K. conceived the project and together with L.P. and J.M.P wrote the manuscript. All authors read and approved the manuscript.

## Acknowledgements

We thank the animal care takers and veterinarians at the animal facility of the Medical University of Vienna for expert help. Dr. Reinhard Grabherr, University of Natural Resources and Life Sciences, Vienna, Austria, kindly provided us with recombinant spike protein. We thank Hans-Christian Theussl, Rubina Koglgruber and Domagoj Cikes at the Institute for Molecular Biotechnology, Vienna, Austria, for their support. We appreciate support by the flow cytometry core facility of the Medical University of Vienna and the Biomedical Sequencing Facility (BSF) jointly run by the Medical University of Vienna and CeMM.

S.K. is supported by the Austrian Science Fund FWF within the special research programs Immunothrombosis (F54-10) and Chromatin Landscapes (F61-04). R.G. received funding by the Austrian Science Fund FWF (ZK57-B28) and P.S. acknowledges funding by the Austrian Science Fund (FWF) (P31113-B30). A.O.R. and H.S. were supported by the FWF (P 34253-B). B.A. was supported by the Austrian Science Fund (FWF) DK W1212. J.M.P. and the research leading to these results has received funding from the T. von Zastrow foundation, the FWF Wittgenstein award (Z 271-B19), the Austrian Academy of Sciences, the Innovative Medicines Initiative 2 Joint Undertaking (JU) under grant agreement No 101005026, and the Canada 150 Research Chairs Program F18-01336 as well as the Canadian Institutes of Health Research COVID-19 grants F20-02343 and F20-02015.

