## Supplementary Material for "ACE2 is the critical *in vivo* receptor for SARS-CoV-2 in a novel COVID-19 mouse model with TNF- and IFNγ-driven immunopathology"

### *Short title:*

Immunity and critical role of ACE2 in murine COVID-19

### Corresponding Author:

Sylvia Knapp, MD, PhD,

Laboratory of Infection Biology, Dept. of Medicine 1, Medical University of Vienna

Waehringer Guertel 18-20, 1090 Vienna, Austria

### Material and Methods

#### *Animal studies*

For passaging, dose-response and time-course experiments, male 10-12 week old BALB/cJ and C57BL/6J mice obtained from Janvier or bred and maintained at the animal facility of the Medical University of Vienna were used. Mice expressing human ACE2 under control of the human *KERATIN-18* (K18) promoter (Tg[K18-Ace2)2$^{Prlmn}$ (McCray et al., 2007) (herein referred to as *K18 huACE2* mice) were obtained from Jackson Labs and bred in the local animal facility. Ace-2 deficient (*Ace2*$^{-/-}$) mice were generated as described (Crackower et al., 2002) and bred at the animal facility of the Institute of Molecular Biotechnology Austria, Vienna. All experiments involving SARS-CoV-2 or its derivatives were performed in Biosafety Level 3 (BSL-3) facilities at the Medical University of Vienna and performed after approval by the institutional review board of the Austrian Ministry of Sciences (BMBWF-2020-0.253.770) and in accordance with the directives of the EU.

#### *Cell lines*

Simian kidney Vero cells were provided by Christoph Steininger (Medical University of Vienna, ATCC CCL81$^{TM}$). To improve *in vitro* SARS-CoV-2 propagation (Matsuyama et al., 2020), Vero cells were transduced as previously described (Machacek et al., 2016) with the retroviral expression vector pBMN-I-GFP carrying human TMPRSS2, amplified from the human lung adenocarcinoma cell line Calu-3 (ATCC HTB-55$^{TM}$, provided by Walter Berger, Medical University of Vienna) and subcloned via the pBluescript KS(-) plasmid. The transduced cell population was subsequently sorted based on GFP expression to >99% purity, creating Vero$^{TMPRSS2}$ cells. Both cell lines were cultured in Dulbecco´s Modified Eagle´s medium (DMEM, Gibco/Thermo Fisher) supplemented with GlutaMAX, 10% fetal calf serum (FCS; Sigma), 1% MEM Non-Essential Amino Acids Solution (Thermo Fisher) and 1% Sodium-Pyruvate (Thermo Fisher) and grown at 37°C at 5% $CO_2$.

#### *SARS-CoV-2 strains, virus propagation and growth curves*

The human SARS-CoV-2 isolate BetaCoV/Munich/BavPat1/2020 (referred to as *BavPat1*) was kindly provided by Christian Drosten, Charité, Berlin, (Rothe et al., 2020), and distributed by the European Virology Archive (Ref-SKU: 026V-03883). *MaVie16* was generated in this study as described. To generate high titre viral stocks, TMPRSS2-overexpressing Vero cells (Vero$^{TMPRSS2}$) were grown on 175 cm$^2$ tissue culture flasks and were infected with *BavPat1* or *maVie16* at a multiplicity of infection (MOI) of 0.1 in Vero cell culture medium (described above) with reduced (2%) FCS content. The inoculum was calculated based on stock concetrations determined using

$TCID_{50}$ assays (described below) assuming that 1*PFU/ml=0,7*TCID50/ml applying the Poisson distribution for the estimation of PFU. Supernatants from infected cells were collected after 48-72h, clarified by centrifugation (3820 G, 15min), and stored in aliquots at -80°C. To determine stock concentrations, $TCID_{50}$ assays were conducted by infecting Vero cells with 150 µl of 10-fold serial dilutions of viral samples (cell supernatant, cell-free lung homogenate), followed by incubation at 37°C. Cytopatic effects were analyzed by microscopy after 72 h and titers were calculated by the method of Reed and Muench (Reed and Muench, 1938). To compare infectivity and viral replication, Vero cells or Caco-2 cells were infected with *BavPat1* or *maVie16* at a multiplicity of infection (MOI) of 0.05 and incubated at 37 °C. After 1 h, the viral inoculum was removed and replaced with medium. At designated time points, supernatants were removed, cleared by centrifugation (500 g, 5 min) and stored at -80°C until further processing for RNA isolation and quantification of genome copies.

#### *Generation of a mouse-adapted SARS-CoV-2 virus, maVie16*

Mouse-adaptation of SARS-CoV-2 and generation of *maVie16* was achieved by serial passaging through lungs of BALB/c mice (16 passages). Briefly, three anesthetized (isoflurane), male 10-12 week old BALB/c mice were infected intranasally (i.n.) with 2x $10^6$ $TCID_{50}$ *BavPat1* in 50 µl. After three days, the mice were euthanized and their lungs were homogenized using a rotor/stator homogenizer in 8-fold volume of DMEM. After removal of aliquots for RNA isolation (see below), the lung homogenates were cleared by centrifugation (3820 G, 6 min), filtered using round bottom tubes with cell-strainer cap (Falcon), pooled and administered i.n. to the next group of anesthetized naïve male BALB/c mice. This process of i.n. infection and harvest was repeated 15 times and weight loss and body temperature (measured with a rectal rodent thermometer; Braintree Scientific) were monitored as readouts of viral pathogenicity. Viral load was quantified by detection of SARS-CoV-2 RNA in the cleared lung homogenate described below. The *maVie16* stock (collected from the lungs of mice infected with passage 15) was prepared in Vero cells as described above.

#### *Mouse model of COVID-19 (mCOVID-19) and tissue collection*

For dose-response and timecourse experiments, male 10-12 week old BALB/c mice and C57BL/6 mice were infected i.n. with 50 µl DMEM containing indicated doses of *maVie16*. Clinical signs of disease (loss of body weight and/or temperature) were monitored at indicated timepoints. Mice were euthanized by cervical dislocation at indicated endpoints for sample collection or whenever they approached 75% of their starting body weight, in accordance with ethical guidlines.

Blood was collected from euthanized animals from the vena cava and transferred to MiniCollect tubes (Greiner) containing $K_3EDTA$. Seventy-five µl of blood per flow cytometry panel were removed, treated for erythrocyte lysis using ACK buffer buffer (150 mM $NH_4Cl$, 10 mM $KHCO_3$, 0.1 mM $Na_2EDTA$, pH 7.2-7.4) and processed for flow cytometry as described below. The remaining blood was centrifuged 10 min at 1700 G, followed by plasma collection and storage at -80°C.

Whole lungs were harvested and weights recorded. The left lung was assigned to histology and fixed in 7.5% buffered formaline. At least 50 µg of the right lung were kept for tissue homogenisation as described above. For total lung RNA, 80 µl of homogenized tissue was lysed in RLT Buffer (Qiagen) containing 2-mercaptoethanol (Sigma-Aldrich) and stored at -80°C until further use. The remaining homogenized lung tissue was centrifuged (3820 G, 6 min) and the cleared homogenate was frozen at -80°C for subsequent viral RNA isolation (see below). To repepare lung single cell suspensions for flow cytometry, the remaining right lungs were minced in gentleMACS tubes with the GentleMACS dissociator (Miltenyi Biotec; program m_lung_01) and digested in 2.5 ml of digestion medium (RPMI medium [Gibco] containing 5% FCS, 250 U/ml collagenase I [GIBCO] and 20 U/ml DNase I [Sigma]) by shaking at 100 rpm for 30 min at 37°C. Digested samples were then homogenized with the GentleMACS dissociator (program m_lung_02) and filtered over a 70 µm cell strainer (BD Biosciences). After centrifugation (500 G, 5 min, 4°C), the cell pellet was treated for 5 min with ACK to lyse erythrocytes. After stopping erythrocyte lysis by addition of PBS 1% bovine serum albumin (BSA; Sigma), cells were passaged over a 40 µm cell strainer (BD Biosciences), and resuspended for subsequent antibody staining (see below).

#### *RNA isolation and gene expression analysis by real-time PCR*

Total RNA was extracted from mouse lung homogenates and viral RNA was extracted from cleared mouse lung homogenates and cell culture supernatants using the following commercially available kits according to the manufacturer´s instructions: For viral RNA extraction, QIAamp Viral RNA Mini Kit (Qiagen) and E.Z.N.A Viral RNA kit (Omega Bio-tek) were used. Quantification of viral RNA was performed by quantitative reverse transcription PCR (RT-qPCR) as described below. For total RNA extraction, the samples were passed through a QIAshredder (Qiagen) to reduce viscosity and subsequently, RNA was isolated using the ROTI Prep RNA Mini kit (Carlroth). After isolation, RNA concentration was measured using a NanoVue Plus Spectrophotometer (GE Healthcare). For cDNA synthesis, 0.5µg RNA from each sample was reverse-transcribed using the qScript cDNA Synthesis Kit (Quantabio). For subsequnt qPCR reactions, PerfeCTa SYBR

Green SuperMix (Quantabio) was used in a total reaction volume of 15ul. TaqMan primer/probes for mouse inflammatory genes were purchased from Microsynth and are listed below. RT-PCR was performed on a StepOnePlus Real Time PCR System (Applied Biosystems) with following conditions: 95°C for 3 min, followed by 45 amplification cycles (15 sec at 95°C) and elongation (1 min at 60°C).

#### *SARS-CoV-2 quantification by real-time PCR*

To generate of a standard curve for quantification of virus genome copy numbers, a 144 bp fragment of the SARS-CoV-2 genome was amplified from *BavPat1* cDNA (generated using the qScript XLT cDNA supermix [Quanta Bio] with RNA purified with the QIAamp Viral RNA Mini Kit) with simultaneous insertion of XhoI (5' end) and BamHI (3' end) restriction site overhangs by PCR (Touchdown from 70°C to 55°C over 35 cycles) using primers CoV-F3_XhoI and CoV-R3_BamHI and Q5 High-Fidelity DNA Polymerase (New England Biolabs). The purified PCR product was then cloned into pIRES2-GFP1 (Clontech) via directional restriction digest and ligation (XhoI and BamHI Fast Digest enzymes from Thermo Fisher; Ligase from New England Biolabs) to generate the plasmid pIRES2-SC2quant. pIRES2-SC2quant was transfected into One Shot TOP10 Chemically Competent *E. coli*, followed by amplification from single cell colonies, isolation of plasmid DNA (using QIAPrep Mini and Midi Prep Kits [Qiagen]) and validation of the construct by sequencing (Microsynth). Based on the molecular weight of the pIRES2-SC2quant plasmid (3344181.19 Da), a $\log^{10}$ dilution series of the plasmid was prepared to contain 0-$10^{10}$ plasmid copies. These dilutions were included on each plate for quantification of viral load in tissue samples by qPCR. The qPCR was performed on a StepOnePlus Real-Time PCR System (Applied Biosystems) using primers CoV-F3 and CoV-R3 with (FAM/TAMRA-labelled) probe CoV-P3 (all at 10 µM and all obtained from Microsynth) and the Ultraplex 1-Step ToughMix ROX (Quantabio), with an initial 10 min 50°C incubation for cDNA synthesis, followed by 40 amplification cycles of denaturation (15 sec at 95°C) and elongation (1 min at 60°C) (Gu et al., 2020).

#### *SARS-CoV-2 sequencing*

For SARS-CoV-2 genome sequencing, viral RNA was processed as described previously (Agerer et al., 2021; Popa et al., 2020). Briefly, viral RNA was reverse-transcribed with Superscript IV reverse transcriptase (Thermo Fisher Scientific) and viral sequences were amplified with modified primer pools (Itokawa et al., 2020). PCR reactions were pooled and processed for high-throughput sequencing. Amplicons were cleaned up with AMPure XP beads (Beckman Coulter) with a 1:1 ratio. Amplicon concentrations and size distribution were assessed with the Qubit Fluorometric

Quantitation system (Life Technologies), and the 2100 Bioanalyzer system (Agilent). Amplicon concentrations were normalized, and sequencing libraries were prepared using the NEBNext Ultra II DNA Library Prep Kit for Illumina (New England Biolabs) according to the manufacturer's instructions. Library concentrations and size distribution were again assessed as indicated previously and pooled into equimolar amounts for sequencing. Sequencing was carried out on the NovaSeq 6000 platform (Illumina) on a SP flow cell with a read length of 2x 250 bp in paired-end mode at the Biomedical Sequencing Facility (BSF) of the Medical University of Vienna and CeMM (Research Center for Molecular Medicine of the Austrian Academy of Sciences). Following demultiplexing, FASTQ files were quality controlled using FastQC (v.0.11.8). Trimming of adapter sequences was performed with BBDUK from the BBtools suite (http://jgi.doe.gov/data-and-tools/bbtools). Overlapping read sequences within a pair were corrected for using BBMERGE function from BBTools. Read pairs were mapped on the combined Hg38 and SARS-CoV-2 genome (GenBank: MN908947.3, RefSeq: NC_045512.2) using the BWA-MEM software package with a minimal seed length of 17 (v0.7.17) (Li and Durbin, 2009). Only reads mapping uniquely to the SARS-CoV-2 viral genome were retained. Primer sequences were removed after mapping by masking with iVar (Grubaugh et al., 2019). From the viral reads BAM (binary alignment map) file, the consensus FASTA file was generated using Samtools (v1.9) (Li et al., 2009), mpileup, Bcftools (v 1.9) (Li et al., 2009), and SEQTK (https://github.com/lh3/seqtk). For calling low-frequency variants, the viral read alignment file was realigned using the Viterbi method provided by LoFreq (v2.1.2) (Wilm et al., 2012). After adding InDel qualities, low-frequency variants were called using LoFreq. Variant filtering was performed with LoFreq and Bcftools (v1.9) (Li, 2011). Only variants with a minimum coverage of 75 reads, a minimum phred value of 90, and indels (insertions and deletions) with an HRUN (homopolymer length on the 3' of the variant) below 4 were considered. Based on control experiments described earlier (Agerer *et al.*, 2021; Popa *et al.*, 2020), all analyses were performed on variants with a minimum alternative frequency of 0.02. Annotations of the variants were performed with SnpEff (v4.3) (Cingolani et al., 2012b) and SnpSift (v4.3) (Cingolani et al., 2012a).

#### *Flow cytometry*

For surface staining, single cell suspensions were treated with TruStain fcX (anti-mouse CD16/32; BioLegend) and Fixable Viability Dye eFluor 780 (eBioscience) according to the manufacturer's instructions. Subsequently, fluorescently-labeled antibodies were added to cells and incubated for 20 min at 4°C. Cells were then washed and fixed for 30 min at room temperature using the Fixation Medium of the Fix and Perm Cell Fixation and Permeabilization Kit (Nordic MUbio). Stained and

fixed cell suspensions were analyzed using an LSRFortessa (BD Biosciences). A flow cytometry gating strategy example (infected mouse lung) can be found in Figure S3. After gating for single/live/CD45$^+$ cells (leukocytes), the following marker combinations have been used to identify the following cell types: lung: alveolar macrophages (AMs): CD11c$^+$/MERTK$^+$; Neutrophils: non-AMs/CD11b$^+$/Ly6g$^+$; Monocytes (Ly6c$^+$ and Ly6c$^-$): non-AMs/non-neutrophils/CD11b$^+$/CD115$^+$; NK cells: Ly6g$^-$ or MERTK$^-$/NK1.1$^+$/CD19$^-$; B cells: Ly6g$^-$ or MERTK$^-$/CD19$^+$/NK1.1$^-$; T helper cells: Ly6g$^-$ or MERTK$^-$/CD19$^-$/NK1.1$^-$/CD3$^+$ or MHCII$^+$/CD4$^+$/CD8$^-$; Cytotoxic T cells: Ly6g$^-$ or MERTK$^-$/CD19$^-$/NK1.1$^-$/CD3$^+$ or MHCII$^+$/CD4$^+$/CD8$^-$; plasmacytoid dendritic cells (pDCs): Ly6g$^-$ or MERTK$^-$/CD19$^-$/NK1.1$^-$/CD3$^+$ or MHCII$^+$/CD4$^-$/CD8$^-$/CD11c$^+$/BST2$^+$; conventional dendritic cells (cDCs): Ly6g$^-$ or MERTK$^-$/CD19$^-$/NK1.1$^-$/CD3$^+$ or MHCII$^+$/CD4$^-$/CD8$^-$/BST2$^-$/CD11c$^+$;
blood: neutrophils: CD3$^-$ or CD19$^-$/CD11b$^+$/Ly6g$^+$; monocytes (Ly6c$^+$ and Ly6c$^-$): CD3$^-$ or CD19$^-$/Ly6g$^-$/CD115$^+$/CD11b$^+$; B cells: CD19$^+$/CD3$^-$; T helper cells: CD19$^-$/CD3$^+$/NK1.1$^-$/CD4$^+$/CD8$^-$; Cytotoxic T cells: CD19$^-$/CD3$^+$/NK1.1$^-$/CD8$^+$/CD4$^-$; NK cells: CD19$^-$/CD3$^-$/NK1.1$^+$; pDCs: CD19$^-$/CD3$^-$/NK1.1$^-$/CD11c$^+$/BST2$^+$;
Data were analyzed using FlowJo software (FlowJo LLC) version 10.7. The fluorescently-labeled anti-mouse antibodies (all obtained from BioLegend unless specifically indicated) listed in the key resources table were used throughout this study.

#### *Plasma cytokine analysis*

Plasma cytokine levels were analyzed using the LegendPlex MU Macrophage/Microglia and Th Cytokine (V02) panels (Biolegend) according to the manufacturer's instructions and using using an LSRFortessa. Before flow cytometry analysis, samples were fixed using Fixation Medium of the Fix and Perm Cell Fixation and Permeabilization Kit (Nordic MUbio).

#### *Analysis of SARS-CoV-2 spike protein-specific antibodies*

SARS-CoV-2 spike-specific antibodies were detected by ELISA based on modified, previously described methodologies (Starkl et al., 2020). Briefly, Nunc MaxiSorp flat-bottom plates (Thermo Fisher) were coated overnight at 4°C with 50 µl of purified recombinant SARS-CoV-2 spike protein ectodomain (Klausberger et al., 2021) (produced in CHO-K1 cells), 2 µg/ml in PBS. After washing 3x with PBS 0.05% Tween-20 (Sigma), plates were blocked by incubation with 100 µl of PBS 1% BSA for at least 2 hours at room temperature. Next, plates were washed 3x, followed by incubation with 50 µl of plasma diluted 1:50, 1:200 and 1:800 (for analysis of IgGs) or 1:40 (for analysis of IgA) for 2 hours at 37°C. After another 3 washing steps, 50 µl biotinylated detection antibodies specific of mouse IgG1 (clone A85-1; BD Pharmingen), IgG2b (clone R12-3; BD Pharmingen) or

IgA (polyclonal; Southern Biotech; all diluted 1:1000 in PBS 1% BSA) were added, followed by incubation for 1 hour at room temperature. Plates were then washed again 3x and incubated with 50 µl of horseradish peroxidase-conjugated streptavidin (BD Pharmingen; 1:500 in PBS 1% BSA) for 20 min at room temperature. Finally, plates were washed 5x and detection was performed using the supersensitive TMB liquid substrate (Sigma) and measurement of (after stopping the reaction by addition of 50 µl 2N $H_2SO_4$) absorbance at 450 nm (620 nm reference) on a Sunrise micropalte reader (Tecan). IgG titers were calculated by plotting the plasma dilution that gave half-maximal signal of a reference plasma (a plasma pool obtained from mice 14 days after infection with *maVie16*).

#### *Histopathological analysis*

Lung samples were fixed in 7.5% formalin, embedded in paraffin, cut into 5µm thick sections, followed by staining with Hematoxylin & Eosin or Toluidine Blue. For staining of viral nucleocapsid protein, embedded samples were deparaffinised by immersion in xylene and rehydrated in graded ethanol. After blocking of endogenous peroxidase with 3% $H_2O_2$ in PBS for 10min at room temperature, tissue sections were first subjected to antigen-retrieval using Antigen Unmasking Solution (Vector Labs) for 10 min and then incubated with TRIS-buffered saline containing 0.01 % Tween (TBST) and 5% goat serum for 10 min at room temperature. Afterwards, slides were incubated overnight at 4°C with SARS Nucleocapsid Protein Antibody (Novus Biologicals) 1:1000 in TBST 5% goat serum. Subsequently, histological slides were washed with TBST and incubated for 30 min with biotinylated goat anti-rabbit IgG (Vector Labs) 1:200 in TBST 5% goat serum at room temperature. After washing with TBST, tissue sections were processed using the Vectastain ABC kit (Vector Labs) and DAB Substrate kit (Vector Labs) according to manufacturer´s instructions. Tissue sections were finally stained with Hematoxylin solution (Mayer´s, Sigma-Aldrich), dehydrated and coverslipped. Histopathological scoring was performed in a blinded fashion by a trained pathologist using the following parameters: alveolar collapse, intraalveolar exsudate, alveolar septal thickening, pneumocyte proliferation, bronchiolar epithelial alteration, bronchitis, peribronchiolar inflammation, interstitial and perivascular infiltrate and interstitial fibrosis (0 = none, 1 = mild/focal/few, 2 = severe/diffuse).

#### *Protein modeling*

A comparative model of mACE2 was created using the Swiss Modeller (Waterhouse et al., 2018), based on the cryoEM structure of hACE2 in complex with BOAT1 (PDB-entry 6m18) (Yan et al., 2020). Sequence identity was at 82 %, and a model with an overal QMEAN score of -1.28 was obtained. The model superimposed to a previous model of the trimeric Spike with hACE2 (https://covid.molssi.org//models/#spike-protein-in-complex-with-human-ace2-ace2-spike-binding), to obtain an initial model of the Spike mACE2 complex. All proteins were fully glycosylated following our previously described protocols (Turupcu and Oostenbrink, 2017). Visualization of the *maVie16* mutations in Spike were subsequently created using the mutagenesis wizzard in Pymol (Schrodinger, 2015).

#### In vivo *treatment with recombinant mACE2*

Recombinant murine soluble (rms) ACE2 (amino acids 1-740) was produced in CHO cells and purified as described (Monteil et al., 2020). To test *in vivo* ACE2 interference, mice received daily intranasal treatments with 100 µg rmsACE2 (Apeiron Biologics) or the respective dilution of vehicle. The first dose was given as a mix with *maVie16* in 50 µl, the remaining doses were administered in 40 µl endotoxin-free PBS (Gibco) to isoflurane anesthetized animals.

#### In vivo *cytokine depletion*

For *in vivo* depletion of IFNγ and TNF, mice were intraperitoneally injected on days 1 and 3 after *maVie16* infection with 200 µl endotoxin-free PBS (Gibco) containing either 500 µg rat anti-mouse IFNγ (clone XMG1.2) and 500 µg rat anti-mouse TNF (clone XT22) or 500 µg rat IgG1 isotype control (clone GL113; specific for ß-galactosidase) obtained from Polpharma Biologics.

#### *Statistical analysis*

Statistical analysis was performed using GraphPad Prism 9.1 (GraphPad Software). Details regarding statistical analyses of experiments can be found in the respective figure legends. *P values* ≤ 0.05 were considered statistically significant.

### Reagents and Resources

| REAGENT or RESOURCE | SOURCE | IDENTIFIER |
|---|---|---|
| **Antibodies** | | |
| TruStain FcX (anti-mouse CD16/32) | BioLegend | Cat# 101320; RRID:AB_1574975 |
| eBioscience Fixable Viability Dye eFluor 780 | ThermoFisher | Cat# 65-0865-14 |
| Brilliant Violet 510 anti-mouse CD45 | BioLegend | Cat# 103138; RRID:AB_2563061 |
| PE anti-mouse CD45 | BioLegend | Cat# 103106; RRID:AB_312971 |
| PE/dazzle 594 anti-mouse CD45.2 | BioLegend | Cat# 109845; RRID:AB_2564176 |
| PE anti-mouse CD45R | BioLegend | Cat# 103208; RRID:AB_312993 |
| PE anti-mouse F4/80 | BioLegend | Cat# 123110; RRID:AB_893486 |
| Alexa Fluor 700 anti-mouse CD11b | BioLegend | Cat# 101222; RRID:AB_493705 |
| PE/Cyanine7 anti-mouse CD11b | BioLegend | Cat# 101215; RRID:AB_312798 |
| Brilliant Violet 605 anti-mouse CD11b | BioLegend | Cat# 101237; RRID:AB_11126744 |
| PE anti-mouse CD11b | BioLegend | Cat# 101208; RRID:AB_312791 |
| APC anti-mouse CD11c | BioLegend | Cat# 117310; RRID:AB_313779 |
| PE-Texas Red anti-mouse CD11c | ThermoFisher | Cat# MCD11C17; RRID:AB_10373971 |
| PE anti-mouse CD11c | BD Biosciences | Cat# 557401; RRID:AB_396684 |
| Brilliant Violet 510 anti-mouse Ly-6C | BioLegend | Cat# 128033; RRID:AB_2562351 |
| PE/Cyanine7 anti-mouse Ly-6G | BioLegend | Cat# 127617; RRID:AB_1877262 |
| FITC anti-mouse Ly-6G | BioLegend | Cat# 127605; RRID:AB_1236488 |
| Brilliant Violet 510 anti-mouse Ly-6G | BioLegend | Cat# 127633; RRID:AB_2562937 |
| PE anti-mouse Ly6G | BioLegend | Cat# 127608; RRID:AB_1186099 |
| PerCP-Cy 5.5 Siglec-F | BD Biosciences | Cat# 565526; RRID:AB_2739281 |
| eFluor 450 anti-mouse MHC Class II (I-A/I-E) | ThermoFisher | Cat# 48-5321-82; RRID:AB_1272204 |
| Brilliant Violet 605 anti-mouse CD115 | BioLegend | Cat# 135517; RRID:AB_2562760 |
| FITC anti-mouse CD19 | BioLegend | Cat# 115506; RRID:AB_313641 |
| Brilliant Violet 605 anti-mouse CD19 | BioLegend | Cat# 115540; RRID:AB_2563067 |
| PE anti-mouse CD19 | BioLegend | Cat# 115507; RRID:AB_313642 |
| FITC anti-mouse CD3 | BioLegend | Cat# 100204; RRID:AB_312661 |
| PE/Dazzle 594 anti-mouse CD3 | BioLegend | Cat# 100245; RRID:AB_2565882 |

| | | |
|---|---|---|
| PerCP/Cyanine5.5 anti-mouse CD3 | BioLegend | Cat# 100217; RRID:AB_1595597 |
| eFluor450 anti-mouse CD3 | BioLegend | Cat# 100213; RRID:AB_493644 |
| PE anti-mouse CD3 | BioLegend | Cat# 100205; RRID:AB_312662 |
| FITC anti-mouse CD4 | BioLegend | Cat# 100406; RRID:AB_312691 |
| Alexa Fluor 700 anti-mouse CD4 | BioLegend | Cat# 100429; RRID:AB_493698 |
| PerCP/Cyanine5.5 anti-mouse CD4 | BioLegend | Cat# 100433; RRID:AB_893330 |
| PE anti-mouse CD4 | BioLegend | Cat# 100407; RRID:AB_312692 |
| Alexa Fluor 700 anti-mouse CD8 | BioLegend | Cat# 100729; RRID:AB_493702 |
| Pacific Blue anti-mouse CD8a | BioLegend | Cat# 100728; RRID:AB_493426 |
| PE anti-mouse CD8a | BioLegend | Cat# 100707; RRID:AB_312746 |
| PE anti-mouse TCR beta chain | BioLegend | Cat# 109207; RRID:AB_313430 |
| PE anti-mouse TCR gamma/delta | BioLegend | Cat# 118107; RRID:AB_313831 |
| PerCP/Cyanine 5.5 anti-mouse NK-1.1 | BioLegend | Cat# 108727; RRID:AB_2132706 |
| APC anti-mouse NK-1.1 | BioLegend | Cat# 108709; RRID:AB_313396 |
| *PE anti-mouse FcεRIα* | BioLegend | Cat# 134307; RRID:AB_1626104 |
| PE/Cyanine7 anti-mouse CD117 (c-kit) | BioLegend | Cat# 105813; RRID:AB_313222 |
| Brilliant Violet 421 anti-mouse CD193(CCR3) | BioLegend | Cat# 144517; RRID:AB_2565743 |
| Alexa Fluor 647 anti-mouse CD49b (pan-NK cells) | BioLegend | Cat# 108912; RRID:AB_492880 |
| Alexa Fluor700 anti-mouse CD49b | ThermoFisher | Cat# 56-5971-80; RRID:AB_2574506 |
| PE/Dazzle 594 anti-mouse CD64 (FCγRI) | BioLegend | Cat# 139319; RRID:AB_2566558 |
| Alexa Fluor 700 anti-mouse CD43 | BioLegend | Cat# 143213; RRID:AB_2800660 |
| APC anti-mouse CD44 | BioLegend | Cat# 103012; RRID:AB_312963 |
| Brilliant Violet 510 anti-mouse CD138 | BD Biosciences | Cat# 563192; RRID:AB_2738059 |
| PerCP/Cyanine5.5 anti-mouse CD21/CD35 (CR2/CR1) | BioLegend | Cat# 123416; RRID:AB_1595490 |
| Alexa Fluor 647 anti-mouse CD23 | BD Biosciences | Cat# 562826; RRID:AB_2737821 |
| PE-Cyanine 7anti-mouse CD93 (AA4.1) | ThermoFisher | Cat# 25-5892-82; RRID:AB_469659 |
| eFluor450 anti-mouse CD90.2 (Thy1.2) | BioLegend | Cat# 140305; RRID:AB_10645335 |
| FITC anti-mouse MERTK | BioLegend | Cat# 151504; RRID:AB_2617035 |
| PE anti-mouse MERTK | BioLegend | Cat# 151505; RRID:AB_2617036 |

| | | |
|---|---|---|
| Brilliant Violet 605 anti-mouse CD127 (IL-7α) | BioLegend | Cat# 135025; RRID:AB_2562114 |
| eFluor450 anti-mouse IgM | ThermoFisher | Cat# 48-5890-82; RRID:AB_10671539 |
| FITC anti-mouse IgD | BioLegend | Cat# 405704; RRID:AB_315026 |
| PE anti-mouse IgA | ThermoFisher | Cat# 12-4204-83; RRID:AB_465918 |
| Brilliant Violet 605 anti-mouse CD62L | BioLegend | Cat# 104438; RRID:AB_2563058 |
| APC/Cyanine7 anti-mouse TER-119/Erythroid Cells | BioLegend | Cat# 116223; RRID:AB_2137788 |
| Biotin anti-mouse IgG1 | BD Pharmingen | Cat#553441 |
| Biotin anti-mouse IgG2b | BD Pharmingen | Cat#553393 |
| Biotin anti-mouse IgA | Southern Biotech | Cat#1040-08 |
| Rabbit Polyclonal SARS Nucleocapsid Protein Antibody | Novus Biologicals | Cat# NB100-56576 |
| Biotin goat-anti-rabbit IgG | Vector Labs | Cat# BA-1000 |
| Rat-anti-mouse TNF (neutralizing) | In house | Clone XT22 |
| Rat-anti-mouse IFNg (neutralizing) | In house | Clone XMG 1.2 |
| Rat-anti- ß-galactosidase (IgG1 isotype control) | In house | Clone GL113 |
| **Bacterial and virus strains** | | |
| SARS-CoV-2; strain Germany/BavPat1/2020 | Charité, Berlin, Germany | European Virology Archive # 026V-03883 |
| SARS-CoV-2 *MaVie16* | This study | |
| **Chemicals, Peptides and Recombinant Peptides** | | |
| Collagenase I | GIBCO/Thermo Fisher Scientific | *17018029* |
| Dnase I | Sigma | DN25 |
| Dulbecco´s Modified Eagle Medium with high glucose, GlutaMAX$_{TM}$, 25mM Hepes | Thermo Fisher Scientific | Cat#10564011 |

| | | |
|---|---|---|
| MEM Non-essential amino acids (100x) | Thermo Fisher Scientific | |
| Sodium pyruvate (100 mM) | Thermo Fisher Scientific | |
| Penicillin Streptomycin (10,000 U/mL) | Thermo Fisher Scientific | |
| Fetal Bovine Serum | Sigma | Cat# F9665 |
| RLT Plus | Qiagen | Cat# 1053393 |
| 2-Mercaptoethanol | Sigma-Aldrich | Cat# M3148 |
| Antigen Unmasking Solution | Vector Labs | Cat# H3300-250 |
| Goat Serum | Novus Biologicals | Cat# NBP2-23475 |
| Hematoxylin solution (Mayer´s) | Sigma-Aldrich | Cat# MHS16 |
| Recombinant SARS-CoV-2 spike protein ectodomain | Reingard Grabherr, BOKU Vienna | (Klausberger *et al.*, 2021) |
| mrsACE2 | In house (Apeiron Biologics) | (Monteil *et al.*, 2020) |
| **Critical Commercial Assays** | | |
| ROTI Prep RNA Mini | Carl Roth | Cat# 8485.1 |
| QIAamp Viral RNA Mini Kit | Qiagen | Cat# 52904 |
| E.Z.N.A Viral RNA kit | Omega Bio-tek | Cat# R6874 |
| qScript cDNA Synthesis Kit | Quantabio | Cat# 95047-500 |
| PerfeCTa SYBR Green SuperMix | Quantabio | Cat# 95055 |
| Vectastain ABC kit | Vector Labs | Cat# PK-6100 |
| DAB Subtrate kit | Vector Labs | Cat# SK-4100 |
| LEGENDplex MU Macrophage/Microglia Panel | BioLegend | Cat# 740846 |
| LEGENDplex MU Th Cytokine Panel V02 | BioLegend | Cat# 740741 |

| | | |
|---|---|---|
| **Experimental Models: Cell Lines** | | |
| Vero cells | ATCC | CCL-81 |
| Vero TMPRSS2 cells | This manuscript | |
| Caco-2 cells | ATCC | HTB-37 |
| **Experimental Models: Organisms/Strains** | | |
| Mouse/BALB/cJ | Own colony, Jackson Labs | JAX #000651 |
| Mouse/C57BL/6J | Own colony, Jackson Labs | JAX #000664 |
| Mouse/*K18 hACE2* | Own colony, Jackson Labs | JAX #034860 |
| Mouse/*Ace2*$^{-/y}$ | Own colony | Crackower *et al.*, 2002 |
| **Oligonucleotides** | | |
| CoV-F3_XhoI: CTCGAGTTTCCTGGTGATTCTTCTTCAGGT | This manuscript | |
| CoV-R3_BamHI: CCTAGGTCTGAGAGAGGGTCAAGTGC | This manuscript | |
| CoV-F3: TCCTGGTGATTCTTCTTCAGGT | Microsynth | (Gu *et al.*, 2020) |
| CoV-R3: TCTGAGAGAGGGTCAAGTGC | Microsynth | (Gu *et al.*, 2020) |
| CoV-P3: AGCTGCAGCACCAGCTGTCCA (FAM/TAMRA-labelled) | Microsynth | (Gu *et al.*, 2020) |
| Mouse *adar1*: fwd: GATGACCAGTCTGGAGGTGC; rev: GCAGCAAAGCCATGAGATCG | Microsynth | This manuscript |
| Mouse *eif2ak2*: fwd: AAGTACAAGCGCTGGCAGAA rev:GCACCGGGTTTTGTATCGAC | Microsynth | This manuscript |
| Mouse *ifng*: fwd: ACTGGCAAAAGGATGGTGACA; | Microsynth | This manuscript |

| | | |
|---|---|---|
| rev: TGGACCTGTGGGTTGTTGAC | | |
| Mouse *ifit1:*<br>fwd:<br>CAGCAACCATGGGAGAGAATGCTGA<br>rev: GGCACAGTTGCCCCAGGTCG | Microsynth | This manuscript |
| Mouse *il1b*:<br>fwd: CAAAATACCTGTGGCCTTGG,<br>rev: TACCAGTTGGGGAACTCTGC | Microsynth | This manuscript |
| Mouse *il6*:<br>fwd: CCACGGCCTTCCCTACTTCA;<br>rev: TGCAAGTGCATCGTTGTTC | Microsynth | This manuscript |
| Mouse *il10*:<br>fwd: TGAGGCGCTGTCATCGATTT;<br>rev: CATGGCCTTGTAGACACCTT | Microsynth | This manuscript |
| Mouse *tgfb*:<br>fwd: AGCCCGAAGCGGACTAT;<br>rev: TCCACATGTTGCTCCACACT | Microsynth | This manuscript |
| Mouse *tnf*:<br>fwd: GCGTGGAGCTGAGAGATAACC;<br>rev: GATCCCAAAGTAGACCTGCCC | Microsynth | This manuscript |
| Human TMPRSS2<br>fwd: AACCTGGGCGCCTGGGA<br>rev: ACGTCAAGGACGAAGACCATGTG | Microsynth | This manuscript |
| **Plasmids** | | |
| pIRES2-AcGFP1 | Clontech | Cat# 632435 |
| pIRES2-SC2quant | This study | |
| pBluescript KS(-) | Stratagene | Cat# 212208 |
| pBMN-I-GFP | Addgene | Plasmid 1736 |
| pBMN-TMPRSS2-I-GFP | This study | |
| **Software** | | |
| GraphPad Prism 9.1 | | https://www.graphpad.com |
| FlowJo | Becton, Dickinson and Company | https://www.flowjo.com/ |

### Supplemental Figures & Legends

Fig. S1

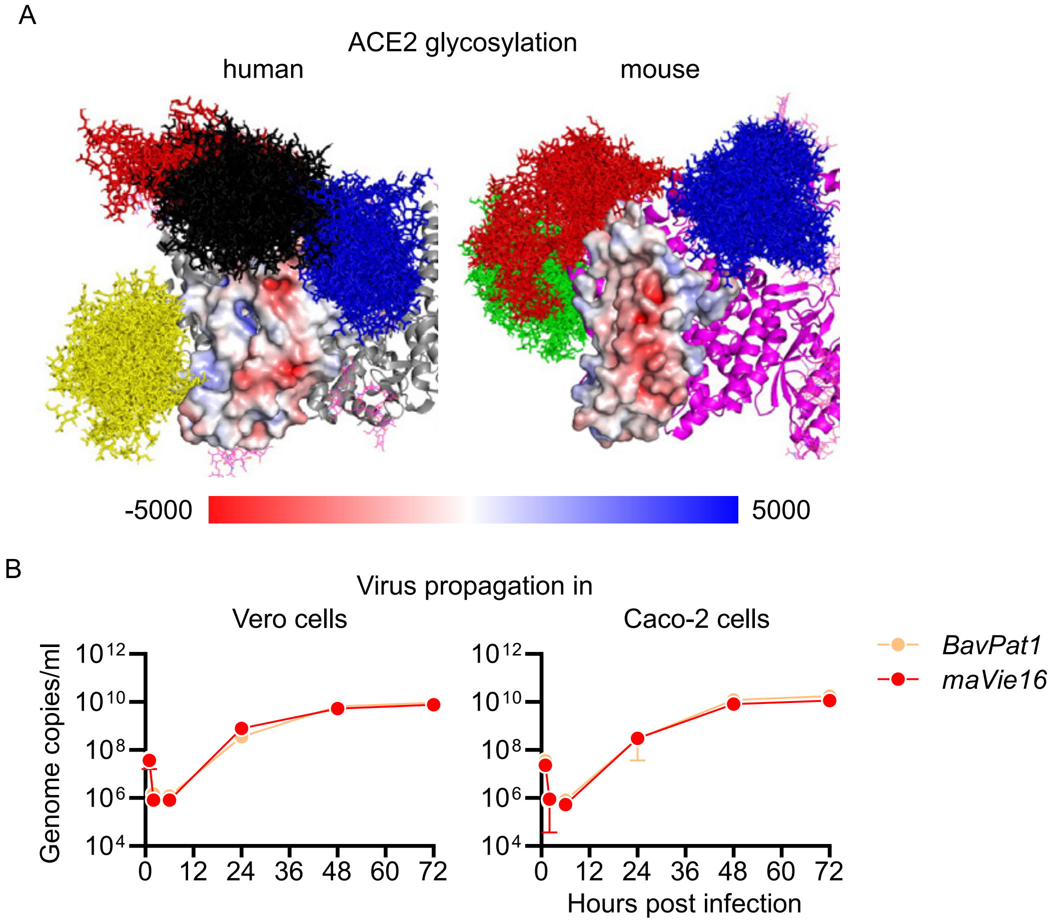
A
ACE2 glycosylation
human
mouse
-5000
5000
B
Virus propagation in
Vero cells
Caco-2 cells
Genome copies/ml
10^12
10^10
10^8
10^6
10^4
0 12 24 36 48 60 72
Hours post infection
BavPat1
maVie16

Figure S1: Mouse *versus* human ACE2 glycosylation and *maVie16 in vitro* proliferation (related to Figure 2)

(A) Surface representation of the binding interface of human (grey cartoon) and mouse (magenta cartoon) ACE2, colored according to the electrostatic potential. Red surface corresponds to a negative potential and blue surface to a positive potential. Note the more distinct pattern of negatively charged areas in the mouse ACE2 protein. A bundle of glycan conformations is shown in sticks for the glycans at N53 (blue), N90 (yellow), N322 (black) and N546 (red) in human ACE2 and at N53 (blue), N536 (green), and N546 (red) in mouse ACE2. (B) SARS-CoV-2 genome copy numbers in Vero and Caco-2 cells at indicated time points after infection with a multiplicity of infection (MOI) 0.5 of *BavPat1* or *maVie16*.

Fig. S2

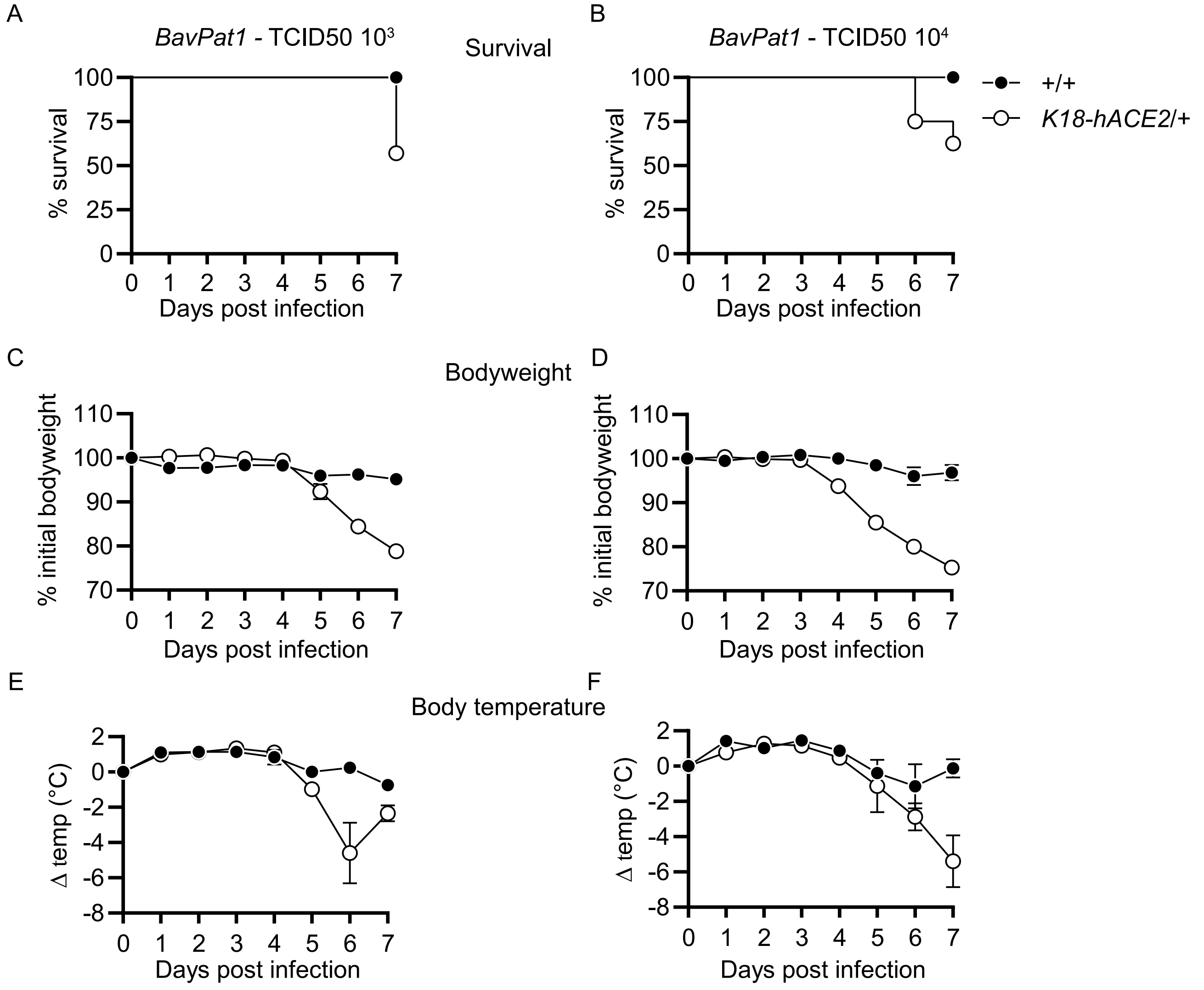

A
B
BavPat1 - TCID50 10³
Survival
BavPat1 - TCID50 10⁴
+/+
K18-hACE2/+
% survival
100
75
50
25
0
0 1 2 3 4 5 6 7
Days post infection
C
Bodyweight
D
% initial bodyweight
110
100
90
80
70
E
Body temperature
F
Δ temp (°C)
2
0
-2
-4
-6
-8

### Figure S2: Disease kinetics of *BavPat1*-infected *K18-hACE2* mice (related to Figure 3)

(A to F) Mice expressing human ACE2 under control of the human *KERATIN-18* promoter (*K18-hACE2*/+) and control animals (+/+) were intranasally inoculated with (A, C and E) $10^3$ or (B, D and F) $10^4$ $TCID_{50}$ of *BavPat1* and monitored over 7 days. (A and B) Survival; (C and D) bodyweight; (E and F) temperature; (A to F) n = 4-7; (E and F) mean +/-SD;

Fig. S3

A

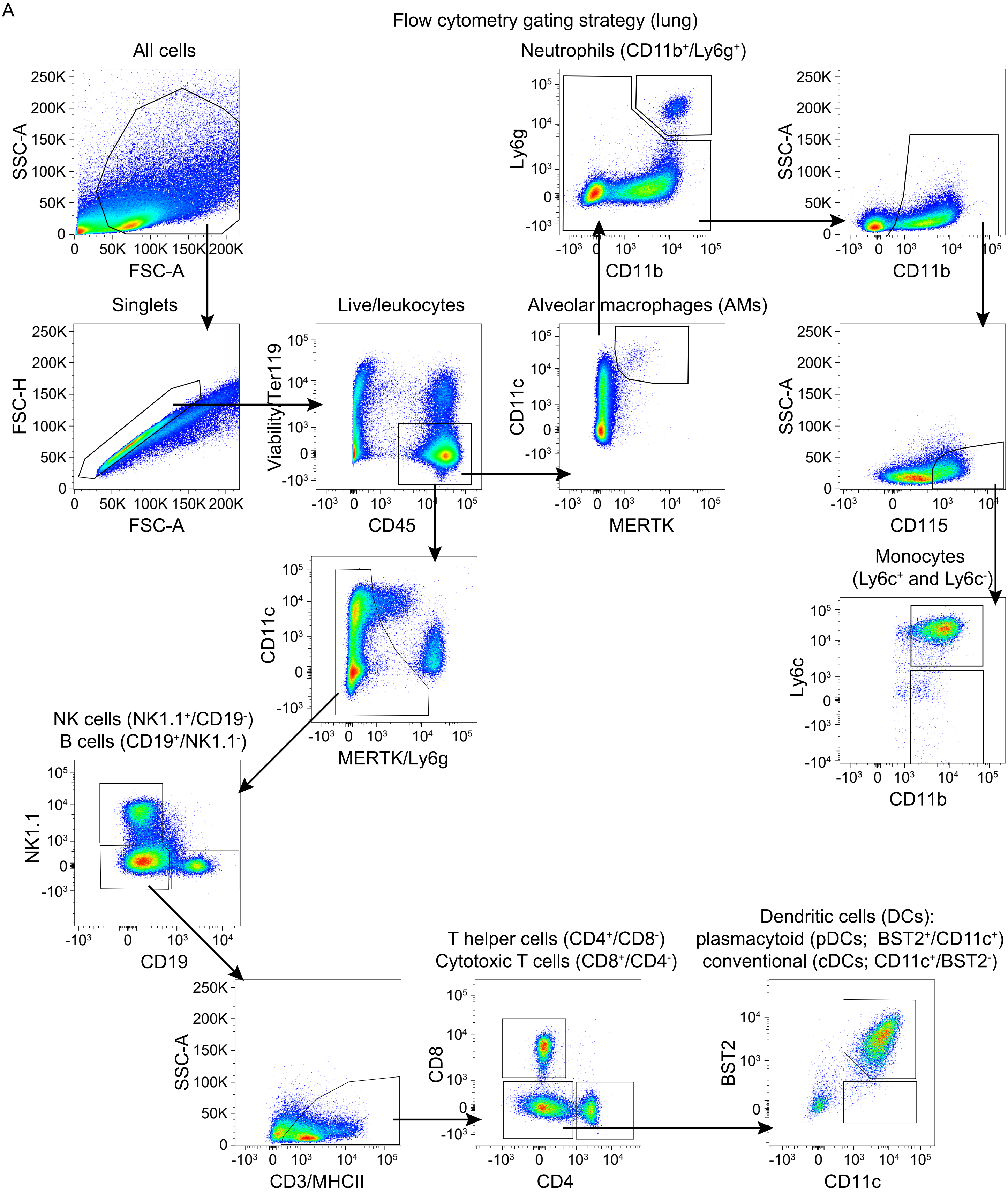

Flow cytometry gating strategy (lung)
All cells
SSC-A
FSC-A
Singlets
FSC-H
FSC-A
Live/leukocytes
Viability/Ter119
CD45
Neutrophils (CD11b⁺/Ly6g⁺)
Ly6g
CD11b
SSC-A
CD11b
Alveolar macrophages (AMs)
CD11c
MERTK
SSC-A
CD115
CD11c
MERTK/Ly6g
Monocytes
(Ly6c⁺ and Ly6c⁻)
Ly6c
CD11b
NK cells (NK1.1⁺/CD19⁻)
B cells (CD19⁺/NK1.1⁻)
NK1.1
CD19
SSC-A
CD3/MHCII
T helper cells (CD4⁺/CD8⁻)
Cytotoxic T cells (CD8⁺/CD4⁻)
CD8
CD4
Dendritic cells (DCs):
plasmacytoid (pDCs; BST2⁺/CD11c⁺)
conventional (cDCs; CD11c⁺/BST2⁻)
BST2
CD11c

### Figure S3: Flow cytometry gating strategy for lung cells (related to Figure 4)

C57BL/6 mice were intranasally infected with PBS (= group 0) or $5x10^5$ $TCID_{50}$ *maVie16* and sacrificed after 2, 5, 7 or 14 days for subsequent analysis. (A) Example of the flow cytometry gating strategy for immune cell populations (shown in Figure 4C and S4B). The sample is derived from an infected mouse 2 days post infection.

Fig. S4

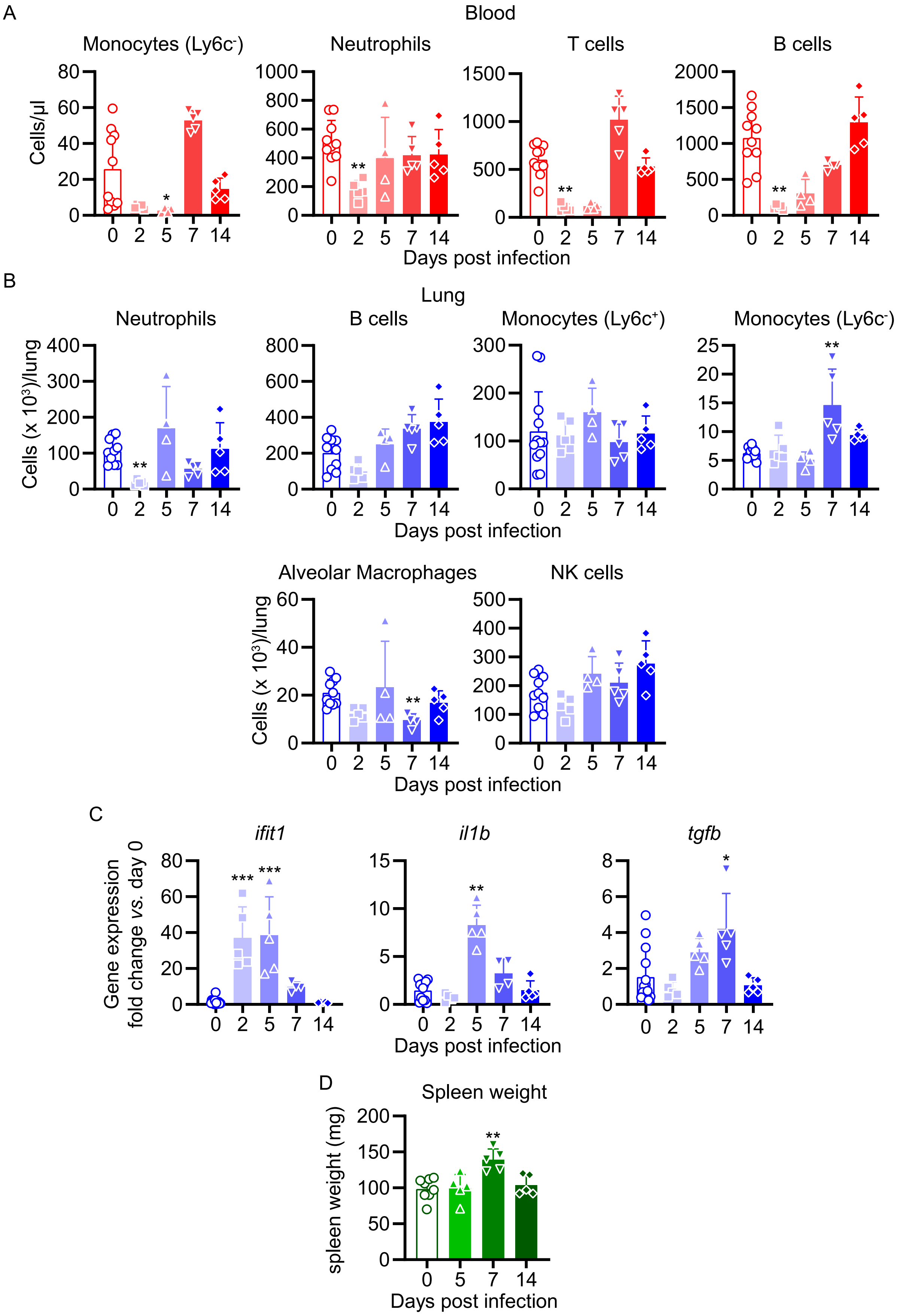

A
Blood
Monocytes (Ly6c-)
Neutrophils
T cells
B cells
Cells/µl
Days post infection
B
Lung
Neutrophils
B cells
Monocytes (Ly6c+)
Monocytes (Ly6c-)
Cells (x 10³)/lung
Days post infection
Alveolar Macrophages
NK cells
Cells (x 10³)/lung
Days post infection
C
ifit1
il1b
tgfb
Gene expression fold change vs. day 0
Days post infection
D
Spleen weight
spleen weight (mg)
Days post infection

### Figure S4: Cellular, transcriptional and spleen weight kinetics of *maVie16*-infected C57BL/6 mice (related to Figure 4)

C57BL/6 mice were intranasally infected with PBS (= group 0) or $5 \times 10^5$ $TCID_{50}$ *maVie16* and sacrificed after 2, 5, 7 or 14 days for subsequent analysis. (A) Flow cytometry analysis of blood cell populations. (B) Flow cytometry analysis of whole lung cell populations (see Figure S3 for gating strategies). (C) Lung tissue expression fold change (compared to group 0 mean; analyzed by real-time PCR) of indicated genes from mice at the respective time points after infection. (D) spleen weight at indicated time points after infection. (A to D): symbols represent individual mice; Kruskal-Wallis test (*vs.* group 0) with Dunn's multiple comparisons test; *$P \leq 0.05$; ** $P \leq 0.01$; *** $P \leq 0.001$.

Fig. S5

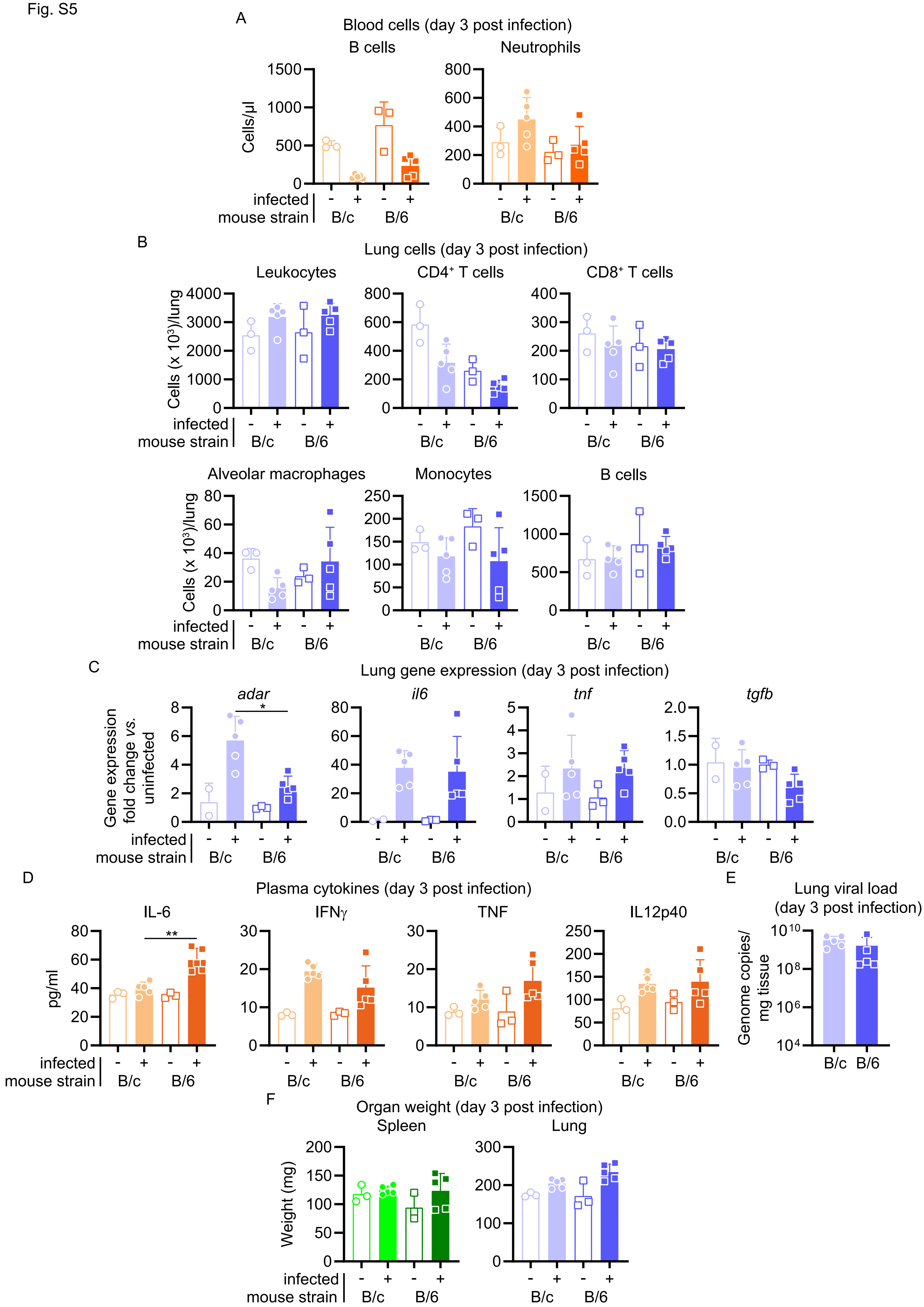

A
Blood cells (day 3 post infection)
B cells
Neutrophils
Cells/µl
infected
mouse strain
B/c
B/6
B
Lung cells (day 3 post infection)
Leukocytes
CD4+ T cells
CD8+ T cells
Cells (x 10³)/lung
Alveolar macrophages
Monocytes
B cells
C
Lung gene expression (day 3 post infection)
adar
il6
tnf
tgfb
Gene expression fold change vs. uninfected
D
Plasma cytokines (day 3 post infection)
IL-6
IFNγ
TNF
IL12p40
pg/ml
E
Lung viral load (day 3 post infection)
Genome copies/ mg tissue
F
Organ weight (day 3 post infection)
Spleen
Lung
Weight (mg)

### Figure S5: Comparison of cellular, transcriptional and organ weight of *maVie16*-infected BALB/c *versus* C57BL/6 mice (related to Figure 6)

BALB/c (B/c) and C57BL/6 (B/6) mice were intranasally inoculated with $10^5$ $TCID_{50}$ *maVie16* (+) or PBS (-). Samples for analyses were collected 3 days after infection. (A) Flow cytometry analysis of blood cell populations. (B) Flow cytometry analysis of indicated lung cell populations in whole tissue. (C) Lung tissue expression fold change (compared to the respective mean of uninfected samples; analyzed by real-time PCR) of indicated genes. (D) plasma levels of indicated cytokines (assessed by multiplex cytokine analysis). (E) Lung tissue virus genome copy numbers (determined by real-time PCR). (F) Weight of spleen and lung. (A to F) mean +SD; symbols represent individual mice; Differences between infected groups were assessed using the Mann-Whitney test; in panels without respective labels, the groups were not significantly different ($P < 0.05$); *$P \leq 0.05$;

Fig. S6

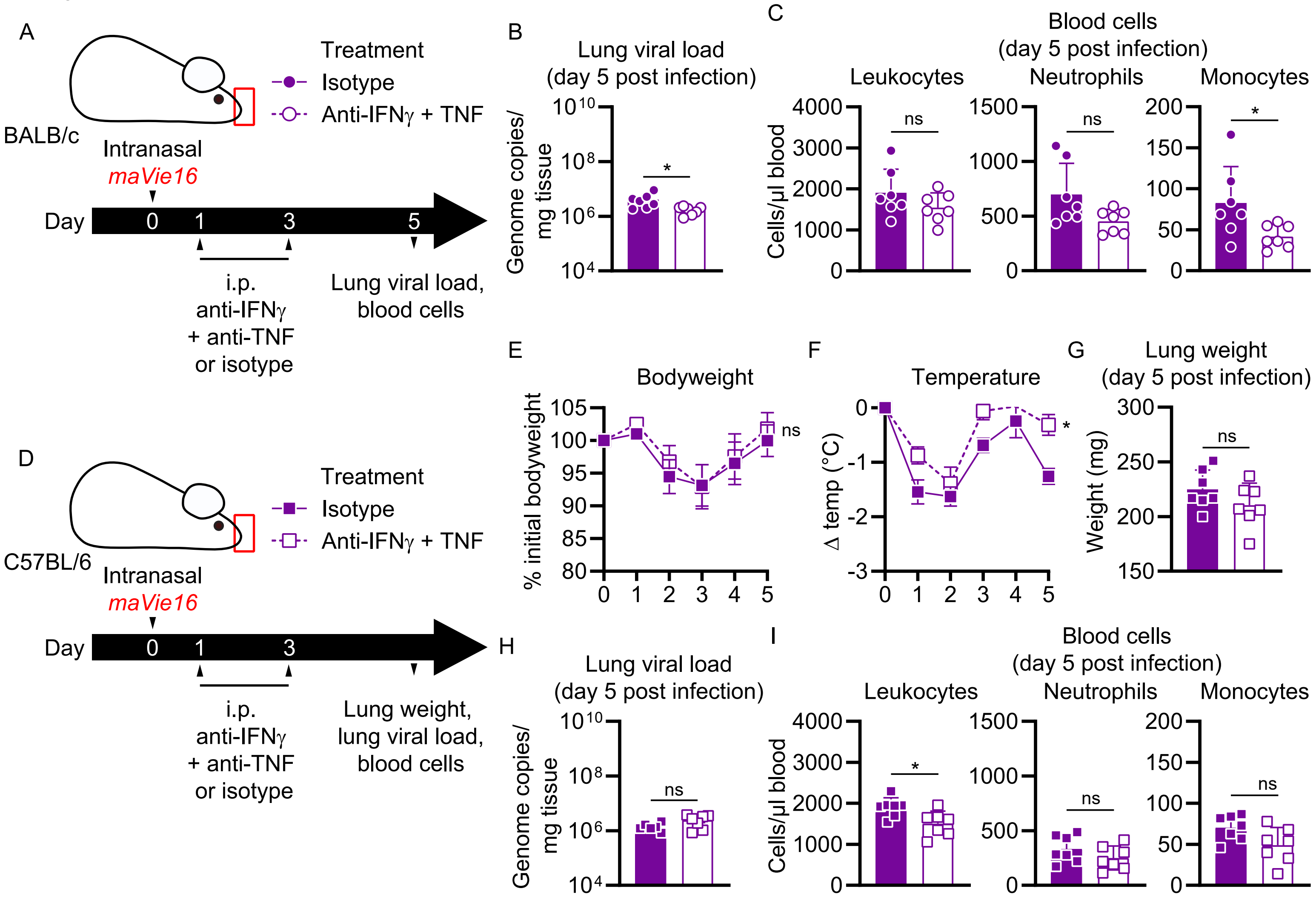
A
BALB/c
Treatment
Isotype
Anti-IFNγ + TNF
Intranasal
maVie16
Day 0 1 3 5
i.p.
anti-IFNγ
+ anti-TNF
or isotype
Lung viral load,
blood cells
B
Lung viral load
(day 5 post infection)
Genome copies/
mg tissue
C
Blood cells
(day 5 post infection)
Leukocytes
Neutrophils
Monocytes
Cells/µl blood
D
C57BL/6
Treatment
Isotype
Anti-IFNγ + TNF
Intranasal
maVie16
Day 0 1 3
i.p.
anti-IFNγ
+ anti-TNF
or isotype
Lung weight,
lung viral load,
blood cells
E
Bodyweight
% initial bodyweight
F
Temperature
Δ temp (°C)
G
Lung weight
(day 5 post infection)
Weight (mg)
H
Lung viral load
(day 5 post infection)
Genome copies/
mg tissue
I
Blood cells
(day 5 post infection)
Leukocytes
Neutrophils
Monocytes
Cells/µl blood
ns
*

Figure S6: Disease parameters of *maVie16*-infected Ace2-deficient mice, and of infected BALB/c and C57BL/6 mice treated with anti-IFNγ and -TNF blocking antibodies (related to Figure 6)

(A) Experimental scheme for B and C. BALB/c mice were infected with $10^5$ $TCID_{50}$ *maVie16* and treated intraperitoneally on day 1 and 3 p.i. with a mix of 500 µg anti-IFNγ and anti-TNF or with isotype control antibody. (B) Lung viral load, and (C) blood cell numbers 5 days p.i.; (D) Experimental scheme for E to I. C57BL/6 mice were infected with 5x $10^5$ $TCID_{50}$ *maVie16* and treated intraperitoneally on day 1 and 3 p.i. with a mix of 500 µg anti-IFNγ and anti-TNF or with isotype control antibody. (E) Bodyweight and (F) temperature kinetics over 5 days after infection. (G) Lung weight, (H) Lung viral load, and (I) blood cell numbers 5 days p.i.; (B, C, G to I) mean +SD; Mann-Whitney test; (E, F) mean +/-SD; 2way ANOVA with Dunnett's multiple comparisons test (*vs.* the respective initial bodyweight or temperature); * $P \leq 0.05$; ns: not significant ($P > 0.05$)

Fig. S7

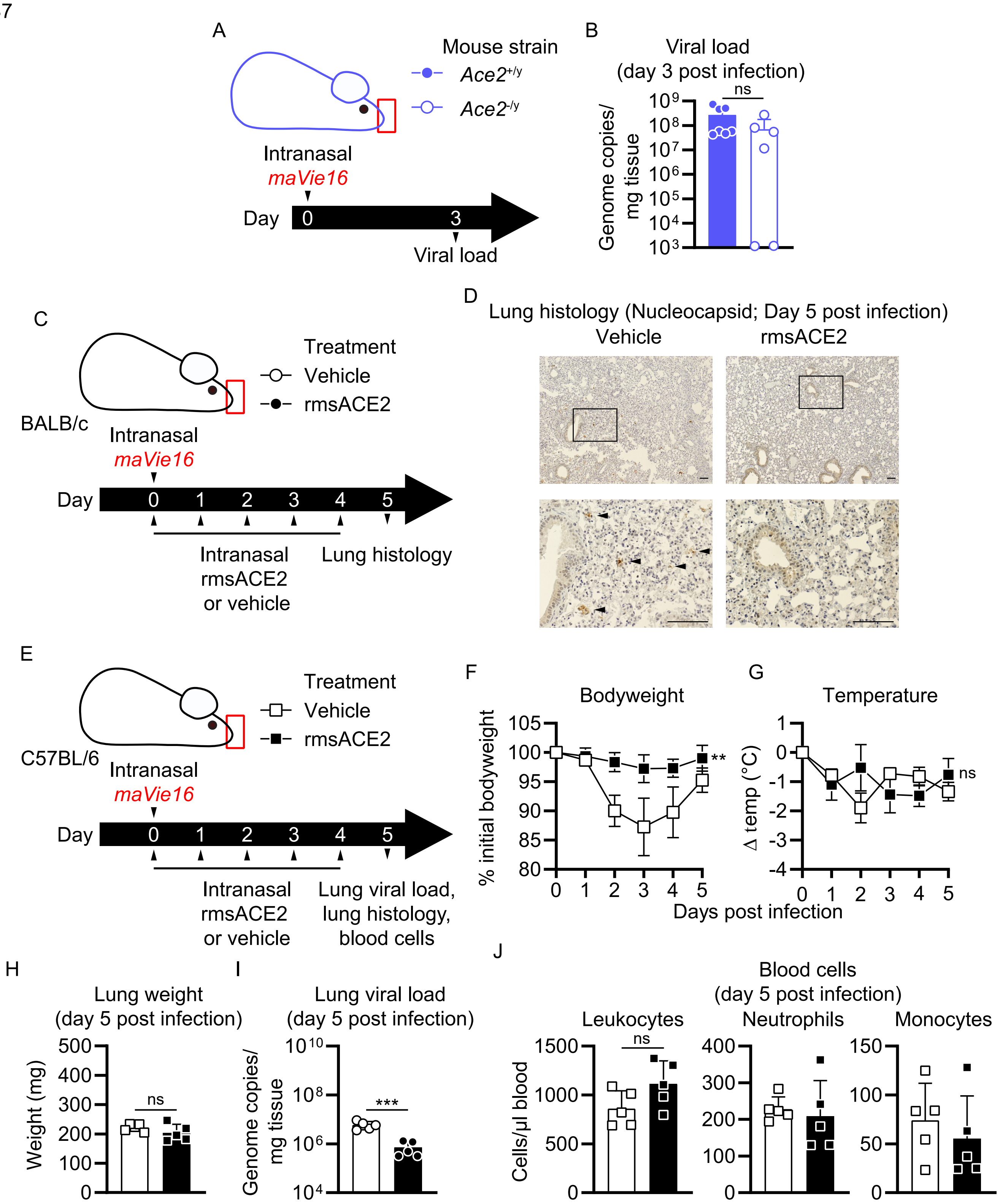

A
Mouse strain
Ace2+/y
Ace2-/y
Intranasal
maVie16
Day
0
3
Viral load
B
Viral load
(day 3 post infection)
ns
Genome copies/
mg tissue
C
Treatment
Vehicle
rmsACE2
BALB/c
Intranasal
maVie16
Day
0 1 2 3 4 5
Intranasal
rmsACE2
or vehicle
Lung histology
D
Lung histology (Nucleocapsid; Day 5 post infection)
Vehicle
rmsACE2
E
Treatment
Vehicle
rmsACE2
C57BL/6
Intranasal
maVie16
Day
0 1 2 3 4 5
Intranasal
rmsACE2
or vehicle
Lung viral load,
lung histology,
blood cells
F
Bodyweight
% initial bodyweight
**
G
Temperature
Δ temp (°C)
ns
Days post infection
H
Lung weight
(day 5 post infection)
Weight (mg)
ns
I
Lung viral load
(day 5 post infection)
Genome copies/
mg tissue
***
J
Blood cells
(day 5 post infection)
Leukocytes
Neutrophils
Monocytes
Cells/μl blood
ns

Figure S7: Disease parameters of *maVie16*-infected Ace2-deficient mice, and of infected BALB/c and C57BL/6 mice treated with recombinant mouse ACE2 (related to Figure 7)

(A) Experimental scheme for B: Male *Ace2*-deficient (*Ace2*$^{-/y}$) or control (*Ace2*$^{+/y}$) mice were infected with 5x $10^5$ $TCID_{50}$ *maVie16*. (B) Lung viral load 3 days post infection (p.i.). (C) Experimental scheme for D. BALB/c mice were infected with $10^5$ $TCID_{50}$ *maVie16* and treated daily intranasally up to day 4 p.i. with 100 µg recombinant murine soluble (rms) ACE2 or vehicle (the first treatment was administered together with virus). (D) lung histology (anti-SARS-CoV-2 nucleocapsid immune-stain) on day 5 after infection; black rectangles in the upper pictures indicate the area magnified in the respective lower row picture; arrow heads indicate infected (nucleocapsid-positive) cells; scale bars represent 100 µm;. (E) Experimental scheme for F to J. C57BL/6 mice were infected with 5x $10^5$ $TCID_{50}$ *maVie16* and treated daily intranasally up to day 4 p.i. with 100 µg recombinant murine soluble (rms) ACE2 or vehicle (the first treatment was administered together with virus). (F) Bodyweight and (G) temperature kinetics over 5 days after infection. (H) Lung weight, (I) lung viral load, and (J) blood cells on day 5 after infection; (B, H to J) mean +SD; Mann-Whitney test; (F, G) mean +/-SD; 2way ANOVA with Dunnett's multiple comparisons test (*vs.* the respective initial bodyweight or temperature); ** $P \leq 0.01$; *** $P \leq 0.001$; ns: not significant ($P > 0.05$)
